# Ligand-receptor promiscuity enables cellular addressing

**DOI:** 10.1101/2020.12.08.412643

**Authors:** Christina J. Su, Arvind Murugan, James M. Linton, Akshay Yeluri, Justin Bois, Heidi Klumpe, Yaron E. Antebi, Michael B. Elowitz

## Abstract

In multicellular organisms, secreted ligands selectively activate, or “address,” specific target cell populations to control cell fate decision-making and other processes. Key cell-cell communication pathways use multiple promiscuously interacting ligands and receptors, provoking the question of how addressing specificity can emerge from molecular promiscuity. To investigate this issue, we developed a general mathematical modeling framework based on the bone morphogenetic protein (BMP) pathway architecture. We find that promiscuously interacting ligand-receptor systems allow a small number of ligands, acting in combinations, to address a larger number of individual cell types, each defined by its receptor expression profile. Promiscuous systems outperform seemingly more specific one-to-one signaling architectures in addressing capacity. Combinatorial addressing extends to groups of cell types, is robust to receptor expression noise, grows more powerful with increasing receptor multiplicity, and is maximized by specific biochemical parameter relationships. Together, these results identify fundamental design principles governing cell addressing by ligand combinations.

## Introduction

Communication systems such as email enable individuals to address messages to specific recipients and groups of recipients. In biological systems, it is crucial to activate the right cells at the right time. Addressing is essential for targeted cell-cell communication by allowing signals to activate specific cell types or defined groups of cell types. Uncovering how signaling pathways enable different types of addressing is critical for understanding natural developmental programs and predictively controlling pathway activation of target cell types for regenerative medicine or clinical applications (Epstein, 2011; Poon et al., 2016). However, the principles that enable cell-cell communication systems to address biological messages have generally remained unclear.

The simplest conceivable realization of addressing would use specific, one-to-one interactions between ligands and cognate receptors (Figure 1A, left). This architecture is conceptually straightforward, has been implemented synthetically in the SynNotch system (Morsut et al., 2016), and is extendable, as new orthogonal ligand-receptor pairs can provide additional communication channels without disturbing existing ones. Despite the simplicity of a one-to-one addressing system, most natural cell-cell communication systems instead employ an interconnected, many-to-many architecture (Figure 1A, right). Pathways such as bone morphogenetic protein (BMP) (Dudley and Robertson, 1997; Heldin et al., 1997; Massagué, 1998; Mueller and Nickel, 2012; Nickel and Mueller, 2019; Schmierer and Hill, 2007), Wnt (Llimargas and Lawrence, 2001; Wodarz and Nusse, 1998), Notch (Shimizu et al., 2000a, 2000b), Eph-Ephrin (Dai et al., 2014), and fibroblast growth factor (FGF) (Ornitz et al., 1996; Zhang et al., 2006) exhibit promiscuous interactions among their multiple ligand and receptor variants. However, these ligand and receptor combinations may activate similar downstream targets. These observations provoke the questions of whether and how molecular promiscuity enables addressing, and what advantages it could have over the one-to-one architecture.

**Figure 1:**
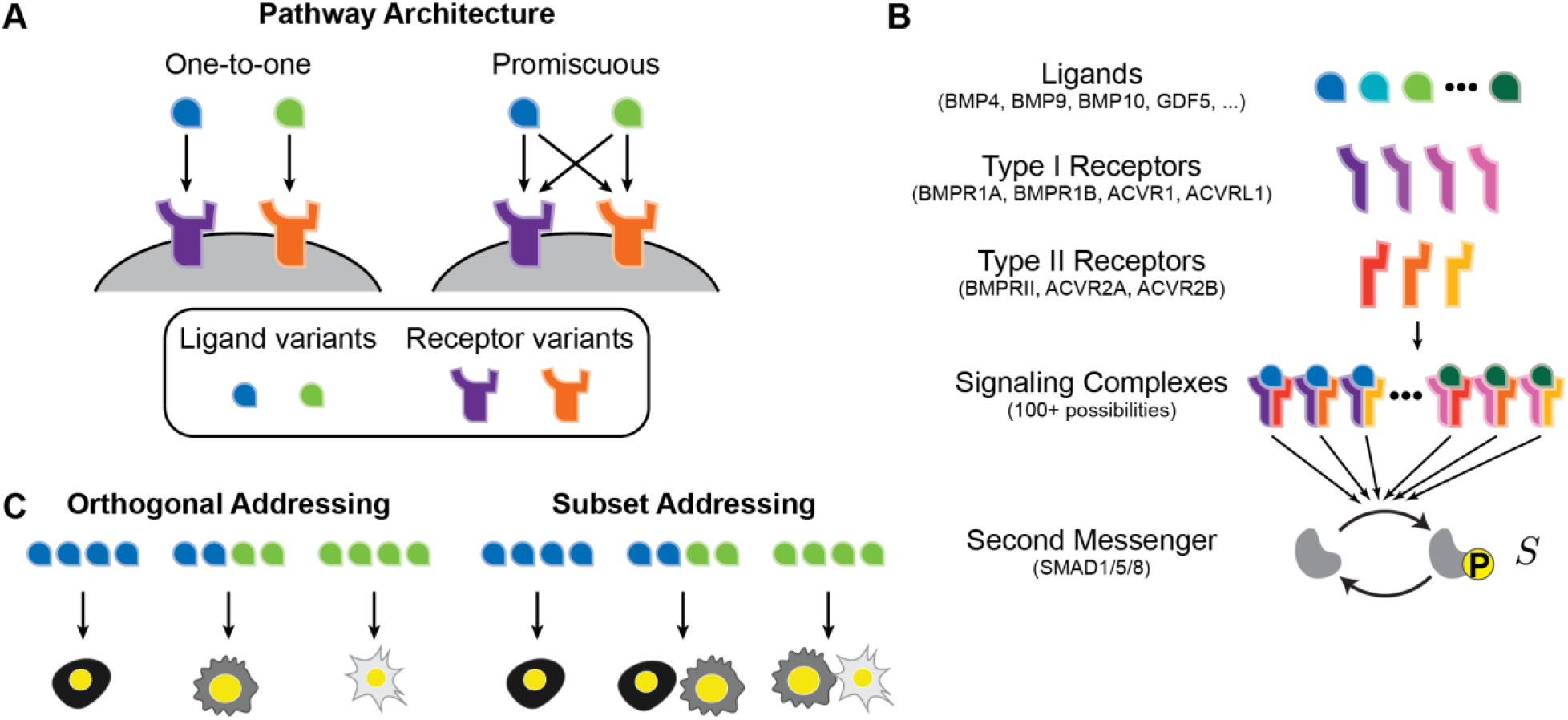
Promiscuous ligand-receptor interactions in the BMP pathway may allow combinatorial addressing. (A) In a one-to-one ligand-receptor architecture, each ligand interacts exclusively with a single receptor (left), while in a promiscuous architecture, ligands interact with multiple receptor variants (right). (B) In this simplified schematic of the BMP pathway, ligands interact combinatorially with type I and type II receptors at the cell membrane to form signaling complexes, which then activate SMAD1/5/8 effector proteins. (C) Signaling pathways could enable different forms of addressing. In orthogonal addressing (left), different combinations of ligands each activate a distinct cell type. More generally, subset addressing (right) could allow activation of different groups of cell types by different ligand combinations.

The BMP pathway provides an ideal system to study these questions. BMP plays diverse roles in most tissues and has demonstrated therapeutic potential (David and Massagué, 2018; Massagué, 2000; Miyazono et al., 2010; Wagner et al., 2010; Wang et al., 2014). In addition, the BMP pathway shows a high degree of promiscuous interactions between its ligands and receptors. In mammals, the pathway comprises more than 10 distinct ligand variants as well as 4 type I and 3 type II receptor variants (Massagué, 2000; Miyazono et al., 2010; Shi and Massagué, 2003). Signaling complexes, comprising a ligand dimer with two type I and two type II receptor subunits, phosphorylate SMAD1/5/8 effectors, which translocate to the nucleus and act as transcription factors to control the expression of target genes (Figure 1B). Individual cells often co-express multiple receptor variants and are exposed to multiple ligands, suggesting that the pathway could function combinatorially (Diez-Roux et al., 2011; Godin et al., 1999; Graham et al., 2014; Kapushesky et al., 2010; Li and Ge, 2011; Liem et al., 1995; Simic and Vukicevic, 2005; Zhang et al., 1998).

Previous observations suggest that BMP ligands could show addressing capacity. For example, during neural tube development, different BMP ligands direct distinct dorsal interneuron identities (Andrews et al., 2017), with each ligand showing specific effects on a subset of interneuron identities but not others. In this way, different ligands appear to address specific progenitors. Moreover, cells also appear to selectively respond to specific ligand combinations in other developmental contexts (Chen et al., 2013; Grassinger et al., 2007; Varley and Maxwell, 1996), and receptor expression patterns can modulate the cellular response (Baur et al., 2000; Lind et al., 1996; Yu et al., 2008).

Recently, mathematical modeling, together with *in vitro* experiments, showed that competitive formation of distinct BMP signaling complexes with different ligands and receptors effectively generates a set of “computations,” in which pathway activity depends on the relative concentrations and identities of multiple ligands (Antebi et al., 2017; Klumpe et al., 2020; Martinez-Hackert et al., 2020). These computations comprise distinct response functions, including additive and ratiometric responses as well as balance and imbalance detection responses that are maximal or minimal, respectively, at defined ligand ratios (Figure S1). Further, the pathway can perform different computations on the same ligands depending on the combinations of receptors expressed by individual cells. In other words, these results suggest the possibility that different ligand combinations could selectively activate, or address, particular cell types based on their receptor expression profiles. Using combinations of ligands to activate specific cell types, promiscuous ligand-receptor interactions may in fact produce additional orthogonal communication channels compared to a one-to-one scheme (Figure 1C, left). In a spatial context, this type of combinatorial addressing could further enable morphogenetic gradients of multiple ligands to activate distinct cell types at specific locations within a tissue.

To understand the principles that govern combinatorial addressing systems, we developed a minimal mathematical model that accounts for promiscuous ligand-receptor interactions, independently representing both the affinities for forming each ligand-receptor signaling complex and their enzymatic activities for activating the pathway. Using a computational optimization approach, we found that promiscuous ligand-receptor interactions generate an extensive repertoire of orthogonal communication channels, exceeding the number possible with the same number of ligands and receptors interacting in a one-to-one fashion (Figure 1C, left). Modest increases in the number of receptor variants substantially increase the number and orthogonality of these addressing channels. Furthermore, the promiscuous architecture allows ligand combinations to address not only individual cell types but also more complex groups of cell types (Figure 1C, right). Finally, using an information theoretic framework, we showed how specific biochemical features, such as anti-correlations between affinity and activity parameters, maximize the information content that can be transmitted through promiscuous ligand-receptor interactions, providing a design principle for building synthetic addressing systems out of promiscuously interacting ligands and receptors.

## Results

### Cell lines show combinatorial addressing *in vitro*

As an initial test of whether the BMP pathway could allow addressing, we analyzed the ability of mixtures of ligands at specific concentrations, or ligand words, to preferentially activate, or address, specific cell types in cell culture, where “cell type” here and throughout the paper refers to a group of cells sharing a common receptor expression profile. (An overview of addressing terminology is provided in Box 1.) To read out pathway activity, we used a transcriptional fluorescent reporter for Smad1/5/8 containing BMP response elements from the Id1 promoter (Korchynskyi and ten Dijke, 2002). We stably integrated the reporter into three cell lines with different receptor expression profiles, then analyzed their responses to a range of BMP ligand combinations by flow cytometry 24 hours after ligand addition (Methods: Addressing of Cell Lines).

Two of the lines were based on a previously characterized epithelial cell line, NAMRU mouse mammary gland (NMuMG) cells, that robustly responds to a variety of BMP ligands (Antebi et al., 2017). In this background, knockdown of ACVR1, which directly interacts with BMP9 (Luo et al., 2010), resulted in a minimal response to BMP9 but a strong response to BMP4, thereby generating a ratiometric response profile (Figure 2A, left). By contrast, knockdown of BMPR2, the major BMP4 receptor (Xia et al., 2007), gave rise to a reduced responsiveness to BMP4 with a strong BMP9 response, producing a complementary ratiometric response (Figure 2A, center). Finally, we also analyzed E14 mouse embryonic stem cells (mESCs) (Figure 2A, right), a different cell type expressing a distinct receptor profile. This line exhibited a synergistic response to BMP4 and BMP9, consistent with previous results (Antebi et al., 2017).

**Figure 2:**
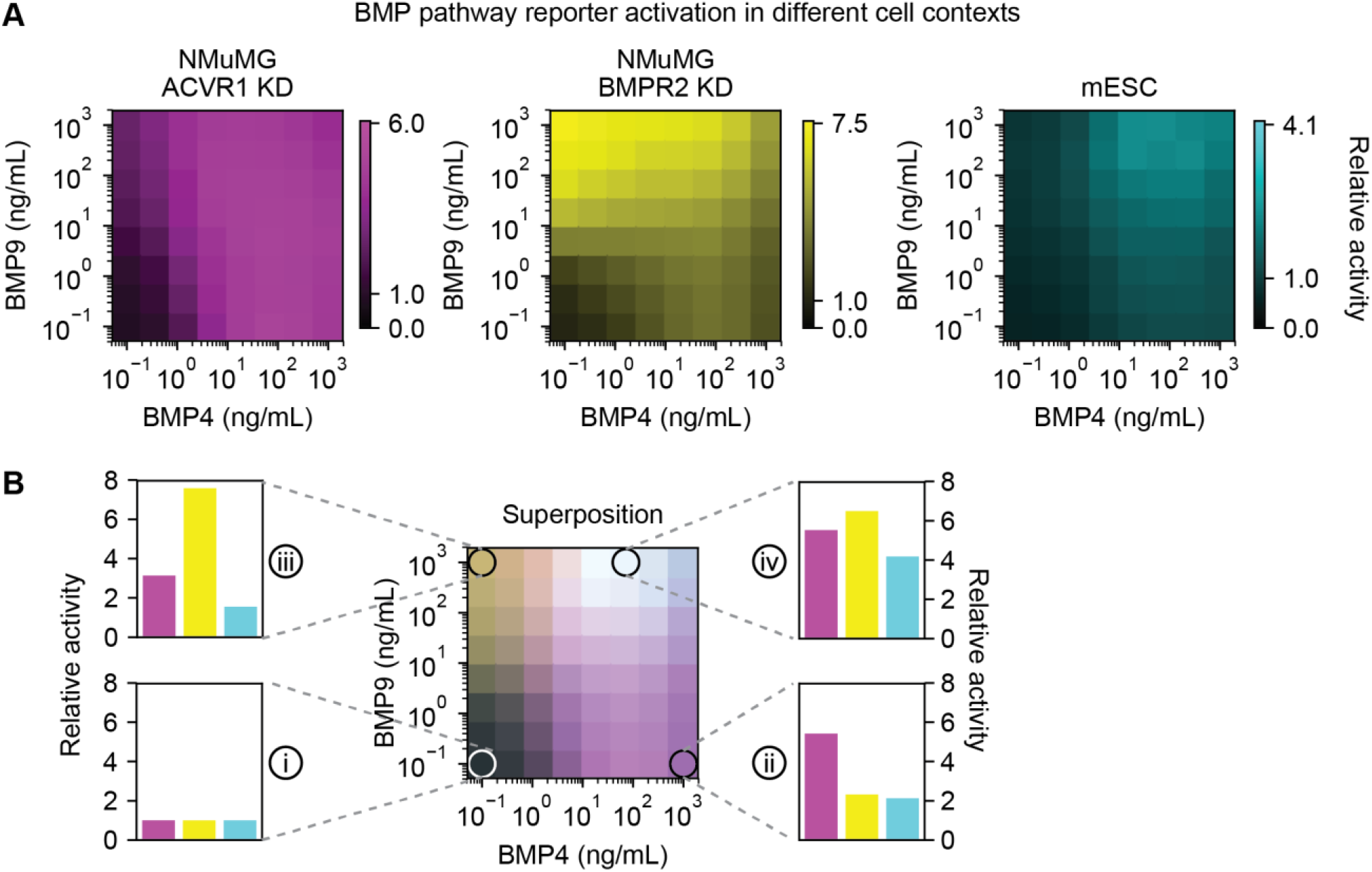
Cell lines preferentially respond to different ligand combinations. (A) Responses of NAMRU mouse mammary gland (NMuMG) cells with siRNA knockdown (KD) of ACVR1 (left), NMuMG cells with siRNA knockdown of BMPR2 (center), or mouse embryonic stem cells (mESCs; right) were measured by flow cytometry of an integrated fluorescent protein reporter (Methods: Addressing of Cell Lines). Each cell line was exposed to a titration of BMP4 and BMP9. Responses are normalized relative to the baseline fluorescence with no added ligand for each cell line. (B) Superimposed responses from (A) reveal that low levels of both ligands (i) generate a basal response. BMP4 alone (ii) preferentially activates NMuMG ACVR1 KD, while BMP9 alone (iii) predominantly activates NMuMG BMPR2 KD. BMP4 and BMP9 together (iv) activate all three cell types to similar levels.

Comparing the responses of these cell lines at different concentrations of BMP4 and BMP9 showed that the two ligands could be used to preferentially activate certain cell types individually or in groups (Figure 2B). For example, BMP4 alone produced 2.3-fold greater response in the NMuMG ACVR1 knockdown cells, compared to the next most responsive cell line. Conversely, BMP9 alone preferentially activated NMuMG BMPR2 knockdown cells, 2.4-fold more strongly than the next most responsive cell line. By contrast, mESCs were strongly activated only when BMP4 and BMP9 were added in combination, a condition that activated all three cell types simultaneously. Overall, distinct combinations (words) of BMP4 and BMP9 preferentially activated three distinct cell type combinations, establishing that the BMP pathway has combinatorial addressing capability. However, the overall potential for addressing remains unknown.

### A minimal model allows analysis of promiscuous BMP ligand-receptor interactions

To explore the addressing capacity of promiscuous ligand-receptor systems, we developed a minimal mathematical model based on the architecture of the BMP pathway (Methods: One-Step Model for Promiscuous Interactions). Briefly, the model describes a set of *n*_*L*_ ligands, *n*_*A*_ type I receptors, and *n*_*B*_ type II receptors. A ligand *L*_*i*_ binds simultaneously to type I and type II receptor subunits *A*_*j*_ and *B*_*k*_ to form an active signaling complex *T*_*ijk*_ (Figure 3A, left). A set of effective interaction strengths, denoted *K*_*ijk*_, represents the strength of binding between a ligand, a type I receptor subunit, and a type II receptor subunit. We further assume that each signaling complex has its own specific activity, denoted *e*_*ijk*_, controlling the rate at which it phosphorylates downstream SMAD effector proteins. The overall activity of the pathway is then the sum of the concentrations of the signaling complexes, each weighted by its own activity parameter. At steady state, this model can be described by one set of equations representing binding and unbinding interactions, a second set of equations representing conservation of total receptor levels, and an expression for total pathway activity, *S* (Figure 3A, right). To solve the model efficiently, we used Equilibrium Toolkit (EQTK), an optimized Python-based numerical solver for biochemical reaction systems (Bois, 2020; Dirks et al., 2007) (Methods: One-Step Model for Promiscuous Interactions). For simplicity, the model neglects some specific features of the natural BMP pathway, including sequential binding of ligands to receptors and the hexameric nature of the full BMP signaling complexes (Massagué, 2000; Shi and Massagué, 2003). These features could enable even greater complexity in pathway behavior beyond that described for this minimal model (Methods: Comparison with Alternative Models).

**Figure 3:**
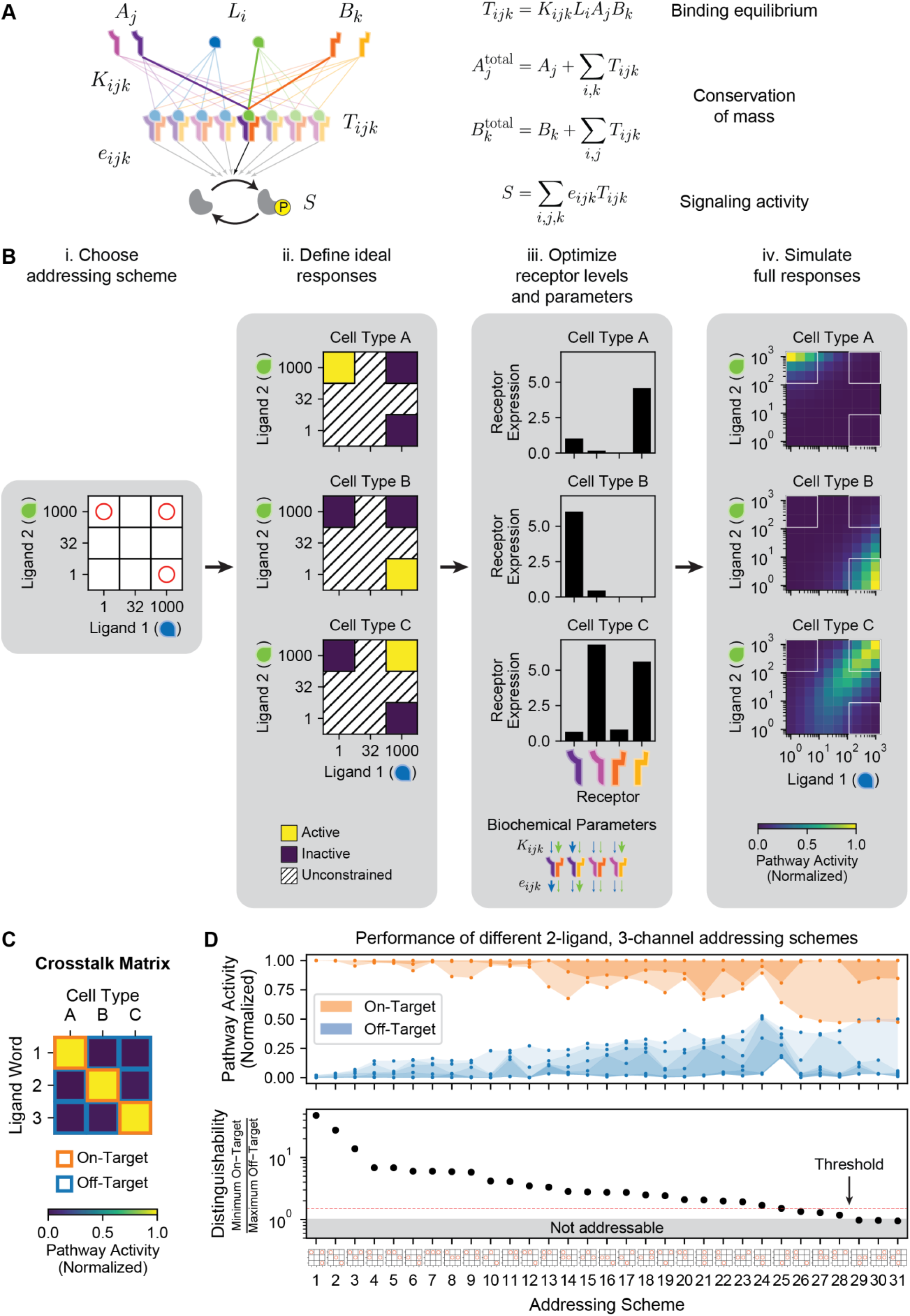
A mathematical model of promiscuous ligand-receptor interactions allows systematic optimization of addressing capabilities. (A) A minimal model of the BMP signaling pathway includes ligand variants (*L*_*i*_; blue and green), which interact with type I receptors (*A*_*j*_; purple and pink), and type II receptors (*B*_*k*_; orange and yellow) to form a combinatorial set of trimeric signaling complexes (*T*_*ijk*_) with varying affinities (*K*_*ijk*_). Active signaling complexes phosphorylate the SMAD effector with varying efficiencies (*e*_*ijk*_). Equations describe the steady-state levels of each component and the total signal *S* (Methods: One-Step Model for Promiscuous Interactions). (B) Optimization systematically identifies potential combinatorial addressing schemes in four steps. i. An orthogonal addressing scheme is specified as orthogonal activation by a set of desired ligand words (red circles). Discretization of ligand space (3×3 grid) enables enumeration of all such addressing schemes. ii. A given orthogonal addressing scheme can be translated into target response functions, in which each cell type is activated by exactly one ligand word (yellow) and not by others (blue). Responses to other ligand words (hatched) are unconstrained. iii. Least-squares optimization identifies a global set of affinity (*K*_*ijk*_) and efficiency (*e*_*ijk*_) parameters, along with a set of receptor expression levels for each cell type, which yield responses similar to the target functions. Upper and lower arrows represent affinity and activity parameters, respectively, for each receptor dimer complexed with each of the two ligands (blue and green arrows). Thin and thick arrows correspond to low and high values, respectively. iv. Responses can be simulated at higher resolution for visualization and further analysis. (C) After optimization, the crosstalk matrix represents the responses of each cell type at the selected ligand words (orthogonal channels). For orthogonal addressing, this matrix should ideally be diagonal, with each ligand word activating only its target cell type (orange border) with no off-target activation (blue border). (D) Best optimization results are shown for all 31 possible 3-channel orthogonal addressing schemes (Methods: Enumeration of Orthogonal Addressing Schemes). (Top) Distributions of on-target (orange) and off-target (blue) activation levels are plotted, representing all elements in the crosstalk matrix. Shaded regions span all activity values. (Bottom) The corresponding distinguishability value for each addressing scheme is shown, along with thresholds at 1 (grey region) and 1.5 (red line). Addressing schemes (x-axis) are shown in order of decreasing distinguishability. See also Figure S2.

### An optimization approach identifies possible addressing schemes

In orthogonal addressing, each ligand word exclusively activates a single cell type, providing one communication channel per cell type. Intuitively, increasing the number of variants of ligand (*n*_*L*_) and receptors (*n*_*A*_ and *n*_*B*_) should expand the number *N* of possible channels by allowing greater diversity of ligand words and cell types. However, it remains unclear whether the number of channels in a promiscuous architecture can exceed the number possible in a one-to-one architecture, how the number of addressable channels grows with increasing ligand and receptor multiplicity, and what biochemical properties enable optimal orthogonal addressing.

To systematically identify parameters that generate orthogonal channels, we used an optimization approach (Figure 3B). We considered discrete ligand concentrations, allowing each ligand to take on one of three logarithmically spaced concentrations, 10^0^ = 1, 10^1.5^ ≈ 32, and 10^3^ = 1000 arbitrary units (AU), reflecting the experimentally observed input dynamic range for BMP signaling (Antebi et al., 2017; Bradford et al., 2019; Hatsell et al., 2015). This discretization defines a finite set of 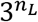 possible ligand words. To identify a system with *N* channels, we chose a subset of *N* ligand words (Figure 3Bi). Each such choice defines an “addressing scheme.” Achieving an addressing scheme requires identifying *N* cell types that are each individually activated by one word (Figure 3Bii). We then used least-squares optimization to identify biochemical parameters (affinities, *K*_*ijk*_, and activities, *e*_*ijk*_) and *N* receptor expression profiles (one for each cell type) that best implement the target addressing scheme (Figure 3Biii; Methods: Optimization of Orthogonal Addressing Schemes). To obtain a more complete view of its functional behavior, we then computed the responses of each cell type on a higher-resolution (10×10) grid of ligand levels (Figure 3Biv).

To quantify the channel structure of the resulting communication system, we computed the crosstalk matrix (Figure 3C), where each row is a ligand word, each column is a cell type, and each value represents the normalized response of that cell type to the corresponding ligand word. Diagonal elements of this matrix represent “on-target” signaling, which ideally approach 1. Off-diagonal elements represent “off-target” signaling, ideally 0. To quantify the addressing specificity, we computed a distinguishability score, defined as the fold difference between the highest off-target activity and the lowest on-target activity (Methods: Distinguishability of Orthogonal Channels).

### Two ligands can orthogonally address five distinct cell types

We first applied this procedure to test whether the promiscuous architecture can achieve more channels than a one-to-one system. Since a one-to-one architecture with two ligands allows for only two orthogonal channels (albeit with perfect specificity), we searched for two ligand systems (*n*_*L*_ = 2) that generate three orthogonal channels in a model with two type I and two type II receptor subunits (*n*_*A*_ = 2 and *n*_*B*_ = 2), reflecting the receptor multiplicity seen in *Drosophila*. We enumerated all 31 possible discrete addressing schemes (Methods: Enumeration of Orthogonal Addressing Schemes), optimized parameters for each scheme, and analyzed the resulting responses (Figure 3D). In 28 of the 31 possible schemes, all on-target activity levels (orange shaded regions) exceeded all off-target activity levels (blue shaded regions), giving rise to orthogonal addressing. While any distinguishability value over 1 achieves addressing, we imposed a stricter threshold of at least 1.5 to ensure better separation of on- and off-target signaling. In fact, 25 schemes showed distinguishability scores above this threshold. Thus, a wide variety of addressing schemes are possible in this minimal system. Among these, the best scheme, based on using each ligand individually as well as a word with both ligands at their maximal level, produced a distinguishability of over 45 (Figure 3D, scheme 1). We note that these results represent a lower bound on the potential addressing capacity and specificity, as global optima are not guaranteed.

Addressing schemes can be realized using a variety of combinations of archetypal response functions previously observed in the BMP signaling pathway (Figure S1) (Antebi et al., 2017). For most addressing schemes, two cell types produced opposite ratiometric responses to the two ligands (Figure S2), with the third cell type exhibiting a variety of responses. The third response types included a balance detector, in which the two ligands combined synergistically activated the pathway more than either ligand alone (Figure S2, e.g. schemes 1 and 2); a nonmonotonic response, in which the pathway was most highly activated at intermediate concentrations of a given ligand (e.g. schemes 3 and 4); a distinct ratiometric response (e.g. schemes 18 and 19); and an additive response to the two ligands (e.g. scheme 17). Thus, the ability of cells to access a variety of multi-ligand response functions with different receptor configurations facilitates addressing.

To extend these results to more channels, we repeated this procedure for schemes of up to eight channels. The eight-channel limit reflects the discretization of ligand concentration space and is not inherent in the system. With the fly-like (*n*_*L*_ = 2, *n*_*A*_ = 2, *n*_*B*_ = 2) model, up to five orthogonal channels could be addressed with at least 1.5-fold distinguishability between on- and off-target conditions (Figures 4A-B). One five-channel scheme achieves 3.6-fold separation between channels using a combination of ratiometric, balance detection, and nonmonotonic responses (Figures 4C and S3A). Additional channels significantly reduced the distinguishability (Figure 4B). Taken together, these results demonstrate that two ligands with promiscuous ligand-receptor interactions can address a larger number of cell types.

**Figure 4:**
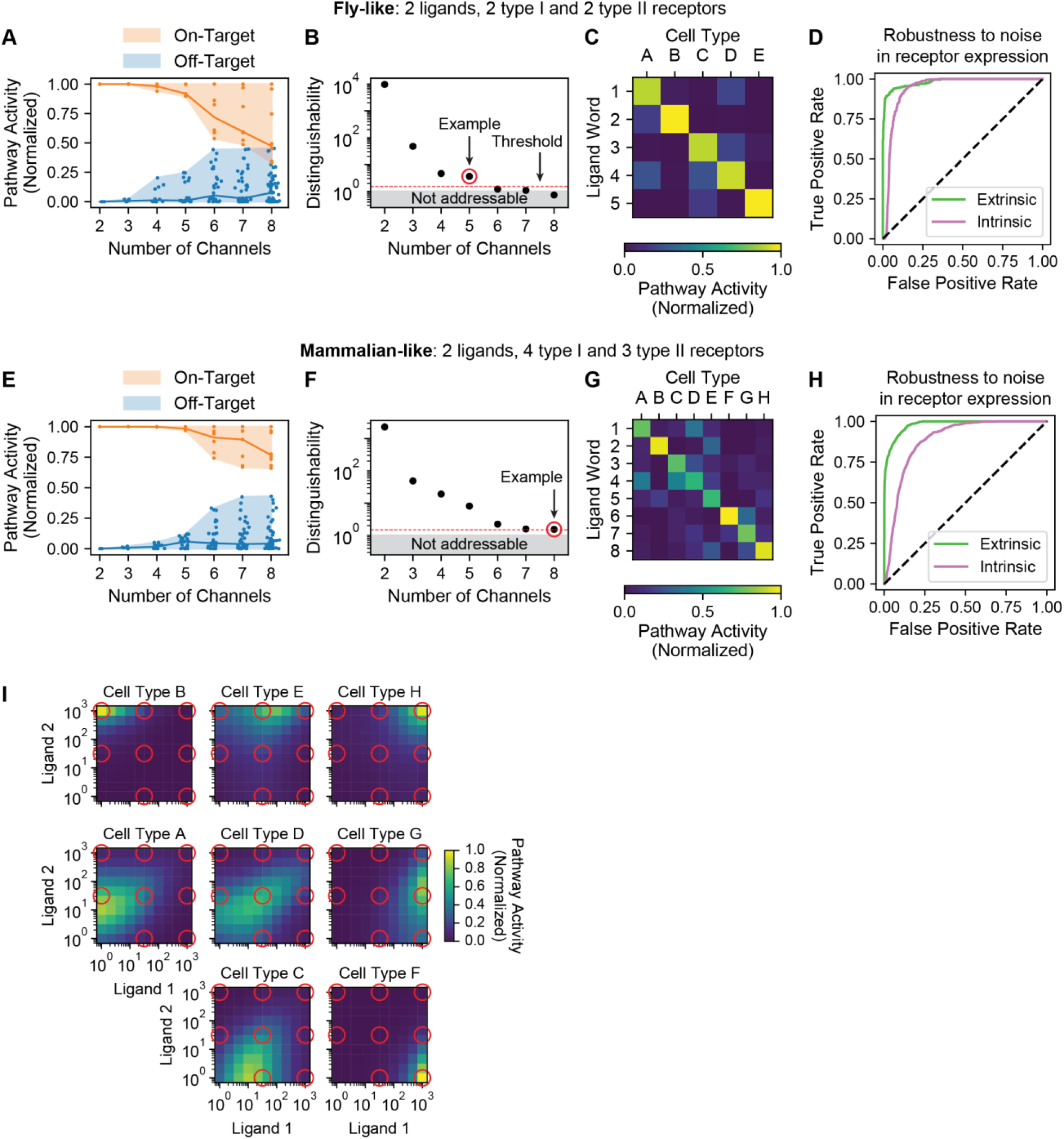
Two ligand variants can independently address eight cell types with high specificity and robustness. (A) In the fly-like model with 2 type I and 2 type II receptor subunits, the pathway activities of each cell type in response to each ligand word are plotted for varying numbers of channels (x-axis), using the optimal parameters for each bandwidth. Shaded regions span full distribution of on-target (orange) and off-target (blue) activities, and lines indicate median values. (B) Distinguishability values are plotted for each number of channels. The 5-channel system (red circle) reflects the highest bandwidth above the threshold of 1.5 (red line). (C) The crosstalk matrix shows the response of each cell type at each ligand word of interest for the 5-channel example circled in (B). Perfect orthogonal specificity would yield a diagonal matrix. (D) Robustness to receptor expression fluctuations was evaluated for the 5-channel system. Optimized receptor expression levels were perturbed in a correlated or uncorrelated way to represent, respectively, extrinsic or intrinsic noise, with a coefficient of variation (CV) of 0.5. The resulting receiver operating characteristic (ROC) curves are plotted and compared to random (black dashed line). (E) The pathway activities for a mammalian-like model with 4 type I and 3 type II receptors are shown, as in (A). (F) Distinguishability values are plotted for the mammalian-like model from (E). The 8-channel system (red circle) reflects the highest bandwidth for a threshold of 1.5 and is further analyzed in (G-I). (G) The crosstalk matrix for the mammalian-like model is shown, as in (C). (H) ROC curves for the 8-channel example are shown, as in (D). (I) The full responses of each cell type are shown for the 8-channel system analyzed in (G-H). Red circles correspond to the 8 ligand words, and cell types are spatially arranged according to the ligand word to which they preferentially respond. For example, the bottom right cell type (cell type F) is orthogonally activated by high levels of ligand 1 only, while the top right cell type (cell type H) would be activated by combining high levels of ligand 1 and 2 together. The bottom left ligand word, with low levels of both ligands, is nonactivating and therefore omitted. See also Figure S3.

### Addressing can occur despite gene expression noise

Stochastic fluctuations, or noise, in gene expression presents a challenge for addressing (Elowitz et al., 2002; Raser and O’Shea, 2005). On the one hand, signaling must be sensitive to receptor expression in order for cell types to have different responses to the same ligand words. On the other hand, if sensitivity is too high, receptor expression noise could disrupt addressing. Here, we asked whether addressing could occur despite correlated (extrinsic noise) and uncorrelated (intrinsic noise) fluctuations in receptor expression. We quantified noise using the coefficient of variation (CV, or s.d./mean) and focused on a physiologically reasonable value of 0.5 (Elowitz et al., 2002; Raj et al., 2006; Suter et al., 2011).

We used two metrics to characterize the extent to which each type of noise degrades addressing. First, we computed receiver operating characteristic (ROC) curves and corresponding area under the curve (AUC) values (Figure 4D; Methods: Analysis of Robustness), which characterize the proportion of on- and off-target cells that are correctly classified (Hanley and McNeil, 1982). (AUC values range from 0.5 for a random system to 1.0 for an ideal system.) Extrinsic and intrinsic noise generated modest reductions in AUC to values of 0.9820 and 0.9400, respectively. Examining the more stringent metric of distinguishability, which is sensitive to incorrect activation of even a single cell type, revealed that intrinsic, but not extrinsic, noise could degrade distinguishability (Figure S3B, left). These results suggest that minimizing intrinsic noise is important for maximizing addressing capacity.

### More receptor variants increase the number of addressable channels

BMP receptor multiplicity has varied during evolution, leading to different numbers of receptor variants in *Drosophila* (2 type I and 2 type II), humans (4 type I and 3 type II), and other species (Massagué, 1998; Newfeld et al., 1999; O’Connor et al., 2006). A model with 4 type I and 3 type II receptor subunit variants, reflecting the multiplicity observed in mammals, significantly outperformed the fly-like model with 2 variants of each receptor subunit (Figure 4E), achieving better specificity at any given number of channels (cf. Figure 4A). In fact, in this model two ligands were able to address as many as eight orthogonal channels with at least 1.5-fold distinguishability between on- and off-target activity (Figure 4F), resulting in a generally diagonal crosstalk matrix (Figure 4G). We note that eight is the maximum number of channels in this three-level ligand discretization scheme; more channels may be possible with higher-resolution grids.

Eight-channel addressing was robust to extrinsic but not intrinsic noise in receptor expression levels, with AUC values of 0.9794 and 0.8853 for extrinsic and intrinsic noise, respectively (Figure 4H). Distinguishability values were generally preserved above 1 for correlated fluctuations of receptor expression but not for uncorrelated noise (Figure S3B, right). This system took advantage of diverse single-cell responses, including ratiometric, balance detection, and nonmonotonic behaviors (Figure 4I). Taken together, these results show that a modest increase in the number of receptor variants generates a substantial expansion in addressing capacity and that the use of multiple distinct response types enables this expanded capacity.

### Promiscuous architectures enable subset addressing

Beyond the addressing of individual cell types, as explored thus far, ligands could in principle generate more complex, multi-cell type response patterns, in which each ligand word activates a specific subset of cell types. In the olfactory system, for example, odorants activate specific subsets of olfactory receptor neurons, giving rise to a combinatorial representation of odors (Hallem and Carlson, 2006; Malnic et al., 1999). Subset addressing systems can be characterized by an “addressing repertoire,” defined as all unique subsets of cell types that can be addressed across all possible ligand words (Figure 5A). Each of these unique subsets represents a channel. For example, a system with 3 cell types that are orthogonally activated would have 3 channels (Figure 5A, top). Higher capacity can occur when some ligand words activate multiple cell types simultaneously (Figure 5A, middle), such as the 4 addressable subsets of the experimental system (cf. Figure 2). The highest bandwidth of 7 addressable subsets occurs when all cell types can be activated in any required combination using some ligand word (Figure 5A, bottom).

**Figure 5:**
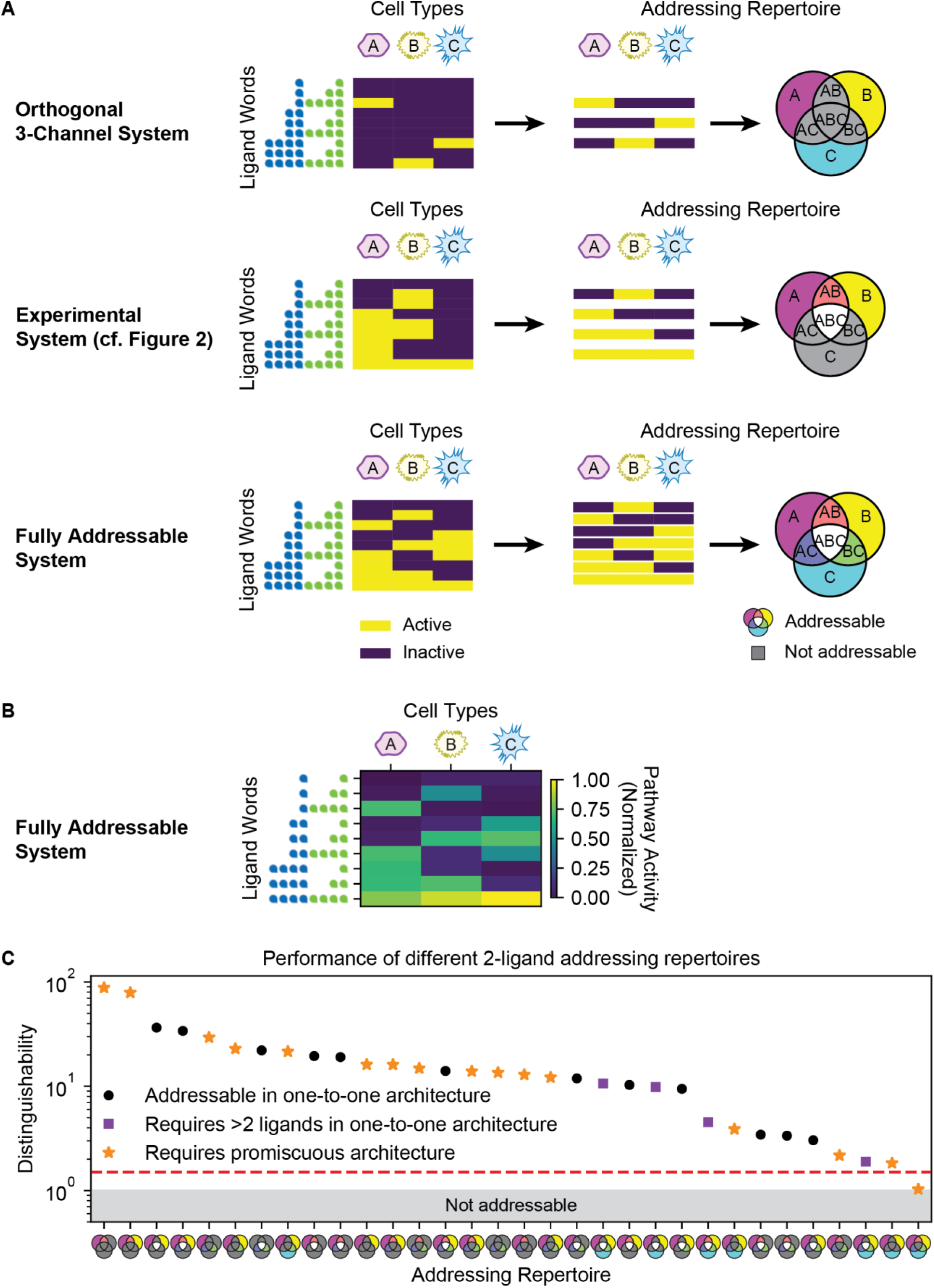
Promiscuous architecture enables diverse addressing repertoires. (A) For three different parameter sets, the responses of three cell types (magenta A, yellow B, and cyan C) to a titration of two ligands (blue and green teardrops) are shown (left). Unique rows reveal the subsets of cell types that can be activated across all ligand words (center). Addressable subsets can also be represented as a Venn diagram (right), where colored regions represent subsets that are activated by at least one ligand combination and grey regions represent subsets that cannot be addressed by any ligand combination. These subsets constitute the “addressing repertoire” of a system. Addressability can vary from purely orthogonal activation (top) to the four subsets shown in the experimental system of Figure 2 (middle) or all possible subsets (bottom). (B) We optimized parameters to achieve the fully addressable system of (A). Simulating the responses of the three cell types to each ligand word confirms that any of the seven possible subsets can be successfully addressed. (C) We generalized the optimization approach to identify parameters achieving each possible addressing repertoire of three cell types in a mammalian-like model with 4 type I and 3 type II receptors. The optimal distinguishability value for each repertoire is plotted. Black circles correspond to addressing repertoires that can be achieved in a one-to-one architecture with two ligands. Purple squares require three or more ligands in a one-to-one architecture, while orange stars indicate addressing repertoires that cannot be achieved in the one-to-one architecture (Methods: Addressing Repertoires).

To determine what addressing repertoires can be achieved in promiscuous ligand-receptor systems, we first considered systems with three cell types, giving rise to seven possible channels. Using the optimization approach for the mammalian-like system of 4 type I and 3 type II receptors, we first sought to identify parameters to achieve this fully addressable system and plotted the corresponding responses of each cell type to each ligand word (Figure 5B), confirming that they do indeed allow for addressing of any of the seven possible channels. We then generalized this approach to all 32 possible addressing repertoires and successfully identified parameter sets that generated 31 with distinguishability values greater than 1.5 (Figure 5C; Methods: Addressing Repertoires). Even the worst repertoire still exhibited distinguishability value greater than 1. Thus, just two ligand variants can generate a broad variety of addressing repertoires.

Achieving such a broad set of addressing repertoires requires promiscuous ligand-receptor interactions. A one-to-one model with two ligands can at most implement two orthogonal channels, and thus would require additional ligands to produce many of the addressing repertoires (Figure 5C, purple squares). Further, half of the addressing repertoires achievable in a promiscuous architecture cannot occur, even theoretically, in a one-to-one architecture with any number of ligands (Figure 5C, orange stars; Methods: Addressing Repertoires). Taken together, these results demonstrate that the promiscuous ligand-receptor architecture allows for an astonishing diversity of addressing repertoires.

### Response function diversity increases addressability

The values of key biochemical parameters – affinities and activities – ultimately determine the addressing bandwidth of a promiscuous ligand-receptor system. What is the distribution of addressing bandwidth across different parameter sets? Are there design rules that allow tuning of those values, in absolute or relative terms, to optimize addressing? Information theory provides a natural framework to answer these questions (Huntley et al., 2016; Itzkovitz et al., 2006). More specifically, the concept of mutual information can be used to quantify the addressing power of a promiscuous ligand-receptor system without assuming any particular choice of ligand words or cell types, or any particular mapping between them (Methods: Computation of Mutual Information).

To identify parameter sets that maximize mutual information, we systematically analyzed the diversity of responses across a set of cell types to a set of ligand words for different biochemical parameter sets (Figure 6A). Mutual information measures information communicated by the optimal subset of ligand words to the optimal subset of cell types, allowing the use of comprehensive libraries (Methods: Libraries of Ligand Words and Cell Types). In a fly-like model, we constructed a discrete ligand word library in which each of two ligands takes on one of three concentration values (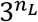 ligand words, or 9); a cell type library, in which each of the 2 type I and 2 type II receptors is expressed at one of two values (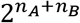 cell types, or 16); and a biochemical parameter library, in which each *K*_*ijk*_ and *e*_*ijk*_ takes on one of two values (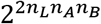 parameter sets, or 65,536). We then simulated the response of each cell type to each ligand word for each biochemical parameter set and computed the mutual information between the sets of ligand words and pathway activity across the library of cell types (Figures 6A-B; Methods: Sampling of Biochemical Parameters). Random, rather than grid-based, sampling of *K*_*ijk*_ and *e*_*ijk*_ produced similar results (Figure S4A). Mutual information values varied broadly across parameter sets, from 0.32 to 1.91 bits, with a median value of 1.36 bits (Figure 6B). By refining our search over biochemical parameters, we were able to identify parameters with values as high as 2.38 bits (Methods: Sampling of Biochemical Parameters).

**Figure 6:**
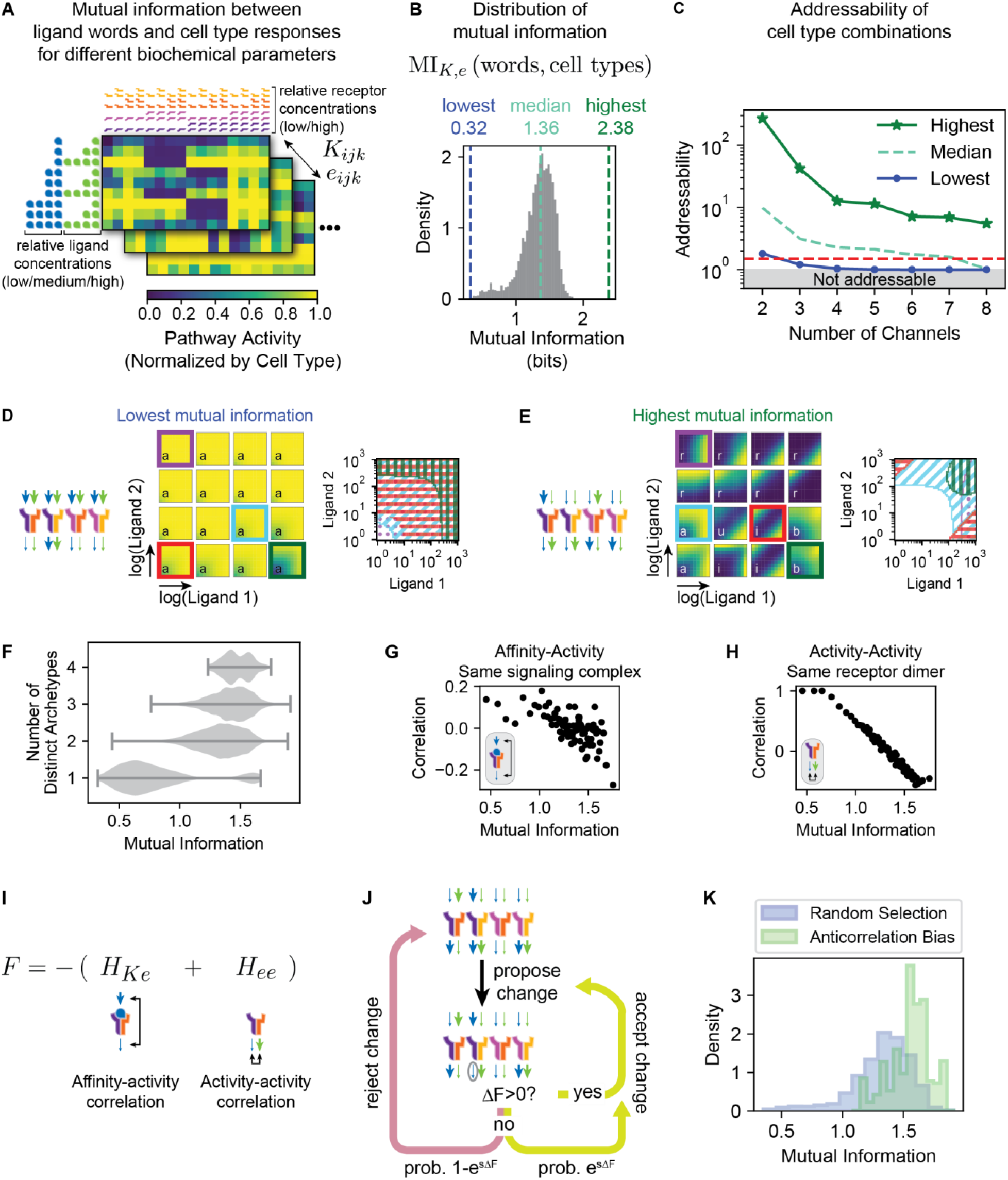
Information theoretic analysis reveals design principles for combinatorial addressing. (A) Mutual information between a comprehensive library of ligand words (rows) and the corresponding activation pattern across a library of cell types (columns) can be computed across a systematic grid-based sampling of the biochemical parameters (*K*, *e*)(matrices). For each row, 1, 2, and 4 ligand symbols indicate low (10^0^), medium (10^1.5^), or high (10^3^) concentrations of the indicated ligand. Similarly, 1 or 2 receptor symbols indicate low (0.1) or high (1) levels of the indicated receptor for each column. (B) The distribution of mutual information across biochemical parameters is shown. Dashed lines indicate the lowest (blue), median (cyan), and highest (green) values. High mutual information indicates that many distinct cell type combinations can be specifically activated by distinct ligand words. (C) The addressability values of activated subsets are shown for different numbers of channels. The addressability reflects the maximal fold difference in the response of at least one cell type when exposed to any two distinct ligand words (Methods: Addressability Metric of Ligand Words). Results are shown for three sets of biochemical parameters generating the lowest, median, and highest mutual information values. (D) The parameter set with the lowest mutual information is represented schematically (left), as in Figure 3Biii. For these parameters, the responses for the library of 16 cell types are shown as a 4×4 grid (center). In each response, the x- and y-axes represent logarithmic titrations of ligands 1 and 2, respectively. All show the same qualitative response of additive (“a”) behavior, differing only in their quantitative sensitivity. Schematically, overlaying four differing responses (highlighted in purple, cyan, red, and green) reveals that different ligand words largely address similar combinations of cell types (right), with relatively few distinct subsets represented. (E) For the parameter set with the highest mutual information (left), the cell types in the library show a variety of response patterns (center): ratiometric (“r”), additive (“a”), imbalance (“i”), and balance (“b”), matching the response archetypes (Figure S1) previously observed experimentally (Antebi et al., 2017). Schematically, overlaying four differing responses (purple, cyan, red, and green) reveals that different ligand words can address many distinct subsets of cell types (right). Note that complexes tend to have opposite values of affinity and activity parameters, as analyzed in (G-H). (F) Violin plots indicate the distribution of mutual information values for systems with different numbers of distinct individual cell response functions (archetypes). Note that greater archetype diversity enriches for high mutual information. (G) Anticorrelations of affinity and activity parameters for the same complex are associated with higher mutual information. We analyzed average properties across bins of 800 parameter sets. To measure the correlation between affinity and activity of complexes, we represented low and high values as −1 and 1 and computed the dot product between *K* and *e* vectors. The average correlation and mutual information across bins are plotted. (H) Parameter sets with high mutual information show anticorrelation in the activities of complexes with the same receptor but different ligands. Analysis was done as in (G). (I) We defined a fitness function *F* that rewards parameter sets exhibiting the anticorrelations observed in (G-H). (J) An evolutionary algorithm identifies parameter sets that maximize *F*. At each iteration, a random parameter value is flipped from low to high or vice versa. Changes that increase *F* are accepted. Changes that decrease *F* are accepted with indicated probability (bottom), which depends on a selection pressure parameter *s*. This process is repeated iteratively (Methods: Evolutionary Algorithm as a Generative Model). (K) An evolutionary algorithm enriches for high mutual information. We ran the algorithm with *s* > 0 to favor anticorrelations or with *s* = 0 to randomly sample parameters. For each case, we randomly initialized 2000 parameter sets and performed 200 iterations. We then evaluated the mutual information for the final value of the parameter set and visualized the resulting distributions. Random selection (*s* = 0, blue) led to a similar distribution of values as the systematically sampled parameter sets (cf. Figure 6B), while favoring anticorrelations (*s* > 0, green) resulted in an overall increase in mutual information. See also Figure S4.

To assess whether mutual information correlates with addressing, we defined an addressability metric, which quantifies how strongly activation patterns differ for different ligand words without requiring specific targeted profiles (Methods: Addressability Metric of Ligand Words). For every pair of ligand words, we identify the largest fold difference of activation levels across all cell types. This value is high when two ligand words induce distinct responses in at least one cell type. The addressability metric is then defined as the lowest such value across all ligand word pairs. We calculate this value for a given number of channels *N* by taking the best choice of all possible subsets of *N* ligand words. Using this metric, we analyzed addressability for systems with low, intermediate, and high mutual information (Figure 6C). For the parameter set of highest mutual information (2.38 bits), each of the 8 ligand words activated a distinct cell type combination with over 5.5-fold addressability. The median parameter set (1.36 bits) addressed up to 7 distinct cell combinations at an addressability of 1.6, while the parameter set with lowest mutual information (0.32 bits) addressed only 2 distinct cell combinations with addressability of at least 1.5 (Figure 6C). Overall, a 1-bit difference in mutual information can increase addressing specificity as well as bandwidth, enabling diverse responses to different ligand words.

We next wanted to understand how high addressing bandwidth arises from the individual response functions of each cell type for the parameter sets with the lowest and highest mutual information (Methods: Analysis of Archetypal Responses). The parameter set with the lowest mutual information generated a homogeneous spectrum of responses across all cell types (Figure 6D). These responses predominantly varied quantitatively in their sensitivity to ligand. By failing to fully exploit the two-dimensional nature of ligand concentration space, this parameter set exhibited limited addressing potential. By contrast, the parameter set with the highest mutual information generated a broad diversity of ligand response functions across the cell types, reproducing the experimentally observed ratiometric, additive, imbalance detection, and balance detection “archetypal” functions (Figure 6E) (Antebi et al., 2017). By generating diverse two-dimensional response functions, this parameter set allows each ligand word to activate a distinctive combination of cell types. In fact, such a correlation between the diversity of response functions and mutual information is seen across the full library of parameter sets (Figure 6F).

### Affinity-activity relationships control addressing bandwidth

We next asked how parameter sets with high mutual information generate the varied response functions associated with addressing. Inspection of the highest mutual information parameter set revealed two striking relationships between binding affinities *K*_*ijk*_ and signaling efficiencies *e*_*ijk*_ (Figure 6E). First, complexes that formed with strong affinity (large *K*_*ijk*_) often had low signaling efficiency (small *e*_*ijk*_). Second, the activity of a given receptor pair strongly depended on the identity of the bound ligand, producing opposite values for *e*_1*jk*_ or *e*_2*jk*_. Systematic analysis of these relationships (Methods: Analysis of Parameter Correlations) revealed their dependence on mutual information and, more specifically, showed anticorrelations between the affinity and activity (*K*_*ijk*_ and *e*_*ijk*_, Figure 6G) and between the activity of complexes with distinct ligands (*e*_1*jk*_ and *e*_2*jk*_, Figure 6H). These results suggest that such anticorrelations could predict high addressing capacity.

To test whether the anticorrelated structure of the parameters is sufficient to produce high mutual information, we developed an evolutionary algorithm that evolves the biochemical parameters to maximize the above anti-correlations (Figures 6I-K). The algorithm starts with an initial parameter set, proposes a random change to one *K*_*ijk*_ or *e*_*ijk*_ value, and accepts that change with probability 1 if the change increases the fitness function *F* and with probability *e*^*sΔF*^ if it does not, where *s* is a parameter that controls the strength of the selection (Methods: Evolutionary Algorithm as a Generative Model). Iteration of this procedure increased mutual information between ligand words and cell types to values comparable to the strongest ones identified in the systematic screen (Figure 6K, cf. Figure 6B).

These results indicate that strong addressing is not confined to a small corner of parameter space. Rather, it is realized to varying degrees across all of *K*, *e* space and enhanced by the parameter anticorrelations identified here. An accompanying experimental analysis of ligand-receptor interactions (Klumpe et al., 2020) suggests that this structure of parameters is present in the natural BMP system. Responses were measured for all pairs of 5 ligands in multiple cell types, which expressed different levels of 2 type I and 3 type II receptors. Fitting a model of receptor competition to these responses predicted anticorrelations between affinity and activity for BMP4, BMP7, BMP9, and BMP10. These four ligands have predicted strong affinity for BMPR1A/BMPR2 receptors but produce low-activity complexes. Similarly, these ligands were predicted to have weak affinity for ACVR1/ACVR2A receptors but produce strong complexes with these receptors. Indeed, analyzing the addressability for each pair of ligands revealed a broad range of addressing capability (Figure S4B). The ligands BMP4, BMP7, and BMP10 together showed the highest addressability, with any pair of these three ligands able to specifically address distinct cell type groups for each ligand combination. Other ligand pairs exhibited lower overall bandwidth but still showed increased specificity at lower bandwidths by an order of magnitude compared to our previously analyzed parameters (cf. Figure 6C). These empirically derived behaviors thus exemplify the anticorrelations that predict high mutual information and addressability.

## Discussion

A fundamental mystery in cell-cell communication is how freely diffusing ligands can precisely target, or address, specific cell types. The promiscuity of ligand-receptor interactions in BMP and other communication pathways makes this question especially perplexing, since it appears to reduce rather than enhance communication specificity. However, promiscuous architectures are employed for specificity in other biological contexts. For example, promiscuous ligand-receptor interactions in the olfactory system enable a limited number of receptors to sense a great diversity of odorants through a combinatorial population code (Duchamp-Viret et al., 1999; Goldman et al., 2005; Hallem and Carlson, 2006; Malnic et al., 1999). Such architectures also appear analogous to simple neural networks, which can compute complex functions of multi-dimensional inputs (Bray, 1995). This computational ability could allow different cell types to respond to different ligand combinations, as observed experimentally (Figure 2) (Antebi et al., 2017; Klumpe et al., 2020).

Our results show that promiscuity indeed allows ligand combinations to address different cell types or groups of cell types with remarkable specificity (Figures 7A-B). While one-to-one architectures can achieve perfect specificity, promiscuous signaling pathways can target more cell types independently (Figure 4) as well as enable greater flexibility in addressing arbitrary subsets of cell types (Figure 5). High addressing capacity can be a robust feature of promiscuous ligand-receptor systems, withstanding correlated noise in receptor expression levels (Figures 4D,H) and emerging across a broad range of biochemical parameter values. A more general mutual information framework identified design principles that maximize addressing capacity (Figure 6). Specifically, these include anticorrelations between the affinity and activity of a given ligand-receptor complex, and anticorrelations between the activities of two ligands interacting with the same receptor dimer. Together, these results show how addressing specificity emerges from molecular promiscuity in a canonical cell-cell communication system.

**Figure 7:**
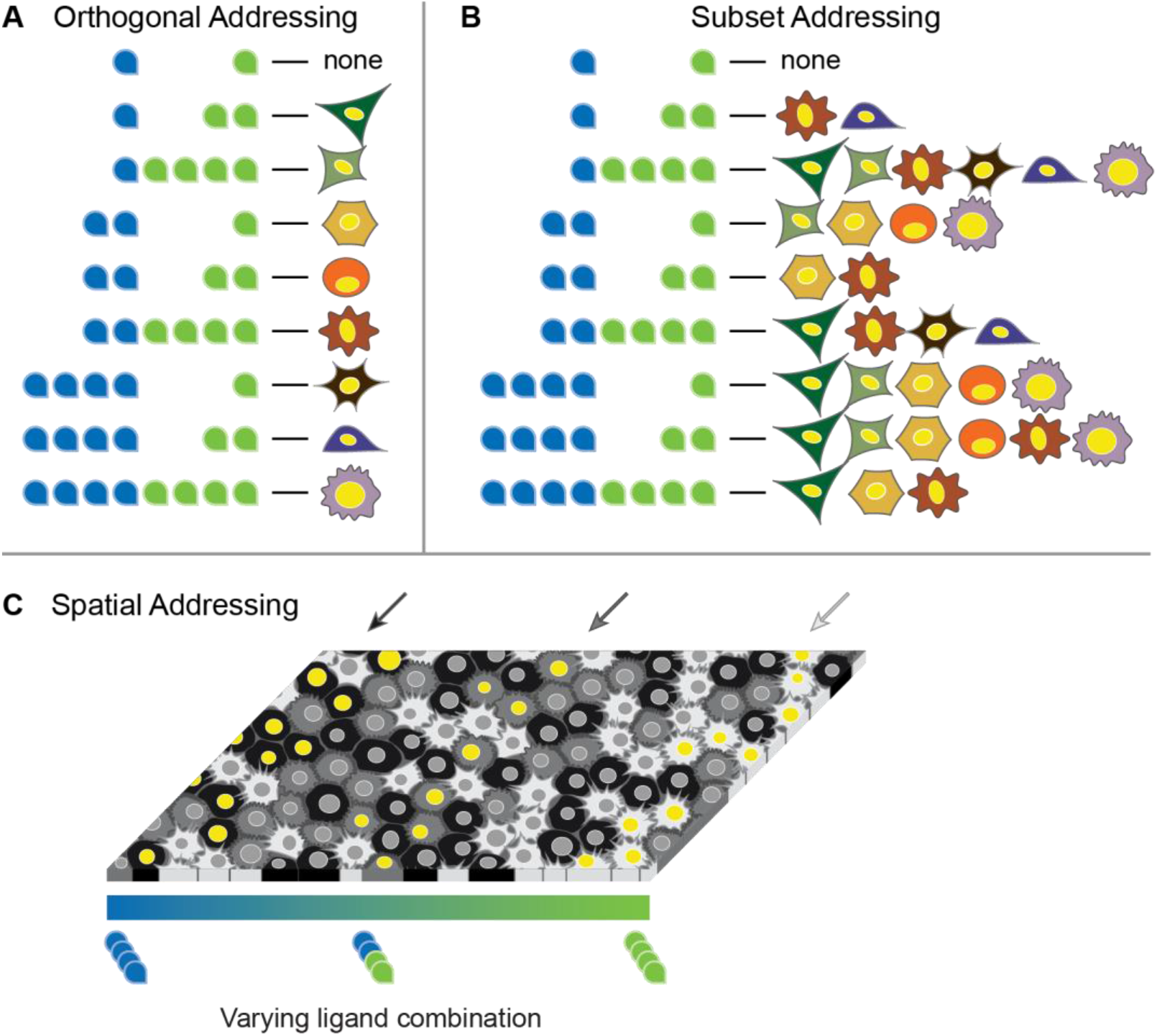
Promiscuous ligand-receptor interactions allow flexible, high-bandwidth addressing. (A) Promiscuous ligand-receptor interactions enable orthogonal addressing, in which individual cell types can be specifically activated using combinations of only two different ligand variants (cf. Figures 4E-I). (B) Promiscuous ligand-receptor interactions enable subset addressing, in which different ligand words address diverse cell type combinations (cf. Figure 6E). (C) Notional schematic showing how two antiparallel morphogen gradients could address different cell types (black, dark grey, and light grey) in specific spatial regions. Yellow nuclei indicate activation. In this example, high levels of blue ligand activate the black cell type (left), the combination of both ligands (blue and green) activates the dark grey cell type (center), and high levels of green ligand activate the light grey cell type (right).

Are the biochemical parameters of the natural BMP pathway compatible with addressing? Systematic analysis of the activity of pairwise ligand combinations showed complex responses to ligand combinations and revealed their dependence on specific receptors (Klumpe et al., 2020). When fit to the same model used here (Figure 3A), these data provide estimated values for *K*_*ijk*_ and *e*_*ijk*_. These values exhibit the types of anticorrelations that favor addressing, suggesting that the BMP pathway may have evolved to facilitate high-capacity addressing.

BMPs function as morphogens, provoking the question of how addressing plays out in a dynamic, spatially extended tissue context. BMP-dependent developmental patterning processes typically use multiple BMP ligands in spatially and temporally overlapping gradients that can be further shaped by shuttling and other extracellular processes. For example, during early *Xenopus* embryo development, an antiparallel gradient of BMP ligands is formed between ventral and dorsal centers (Ben-Zvi et al., 2008; Reversade and De Robertis, 2005). Similarly, overlapping expression patterns of GDF5 and multiple BMP ligands, together with distinct receptor expression patterns, play a key role in activation (and suppression) of BMP signaling in specific cell populations during joint formation (Lyons and Rosen, 2019; Salazar et al., 2016). In such overlapping gradients, addressing could allow different cell types, all with functional BMP pathways, to each selectively respond in distinct regions based on the concentrations of multiple ligands, as shown schematically in Figure 7C. Additionally, temporal changes in receptor expression are common during development (Danesh et al., 2009; Dewulf et al., 1995; Erickson and Shimasaki, 2003; Sanyal et al., 2002). For instance, in neural precursors, *Bmpr1a* is expressed early and ubiquitously; subsequent treatment with BMP2 induces activation of BMPR1A and expression of *Bmpr1b* (Panchision et al., 2001). These different receptor expression states could preferentially respond to different ligand combinations and therefore be addressable. Spatiotemporal addressing could be tested experimentally by genetically modifying the expression of BMP variants in developmental contexts and analyzing the effects on different cell types. *In vitro* reconstitution of multi-ligand gradients could allow a complementary, systematic analysis of spatial addressing (Li et al., 2018).

An increasing amount of receptor expression data is available from cell atlas projects. Together with quantitative measurements of effective biochemical parameters, these data could be used to design ligand combinations that selectively address target cell populations. The ability to design selective targeting would be useful in biomedical applications such as directed differentiation and targeted therapy. For example, recombinant BMP2 has been tested in a variety of therapeutic applications, largely related to promoting bone healing and regrowth. However, there are significant risks, such as ectopic bone formation, respiratory failure, tissue inflammation, and others (Epstein, 2011; Poon et al., 2016). If these complications result from undesired activation of off-target cell types, using a combination of ligands could potentially provide more specific addressing of the appropriate cell type(s). Other potential therapeutic applications for modulators of BMP signaling include cardiac fibrosis, where BMP2 and BMP7 have both shown promise in animal models (Flevaris et al., 2017; Wang et al., 2012); Parkinson disease, where BMP2 and GDF5 both appear to promote survival of dopaminergic neurons (Hegarty et al., 2014; O’Keeffe et al., 2017; O’Sullivan et al., 2010); and cancer, where inhibition of BMP signaling reduces tumor formation in mice (Yokoyama et al., 2017). As the range of clinical applications targeting BMP signaling continues to grow, it will be essential to determine whether combinations of ligands could provide greater specificity than individual ligands.

The principles elucidated here in the context of BMP signaling could apply to other pathways that exhibit promiscuous ligand-receptor interactions, including the broader TGF-β pathway as well as the Wnt, FGF, Eph-Ephrin, and JAK-STAT pathways. The principle of addressing suggests that beyond sensing the concentration of a given set of ligands, these pathways may serve more broadly as computational devices that exploit promiscuous interactions, enabling cells to tune in to specific ligand words and thereby receive information specifically addressed to them.

## Acknowledgments

This work was supported by the Defense Advanced Research Projects Agency (contract HR0011-16-0138), the Gordon and Betty Moore Foundation (grant GBMF2809 to the Caltech Programmable Molecular Technology Initiative), the Human Frontiers Science Program (grant RGP0020), the Institute for Collaborative Biotechnologies (grant W911NF-09-0001 from the U.S. Army Research Office), the National Institutes of Health (NIH) (grants R01 HD075335A and R01 MH116508), and the Paul G. Allen Frontiers Group and Prime Awarding Agency (award UWSC10142). This work does not necessarily reflect the position or policy of the U.S. Government, and no official endorsement should be inferred. C.J.S. is supported by the NIH National Institute of General Medical Sciences (grant T32 GM008042) and a David Geffen Medical Scholarship. H.K. is supported by a National Science Foundation graduate research fellowship (grant DGE-1144469). M.B.E. is a Howard Hughes Medical Institute Investigator.

## Author Contributions

C.J.S., A.M., Y.E.A., and M.B.E. conceived and designed the research. J.M.L., H.K., and Y.E.A. performed experiments. C.J.S., A.M., A.Y., J.B., and Y.E.A. developed mathematical models and performed computational analysis. C.J.S., A.M., Y.E.A., and M.B.E. wrote the manuscript.

## Declaration of Interests

The authors declare no competing interests.

## Box 1: Addressing Terminology

Combinatorial addressing involves mappings between combinations of ligands and responses of cell types defined by their receptor subunit expression. Here, we define some of the terminology introduced in the paper to describe these relationships.

### Ligand word

A set of specific concentration values for each ligand variant in a combination. For example, a concentration of 10 μM for ligand 1 and 100 μM for ligand 2 constitutes a ligand word (10 μM, 100 μM).

### Cell type

A set of specific receptor subunit expression levels. For example, a cell expressing receptor subunits 1 and 3 would represent a different cell type than a cell expressing subunits 1, 2, and 4 or a cell expressing more subunit 1 and less subunit 3.

### Channel

A set of one or more cell types that can be selectively activated (without activating other cell types) by some ligand word. For example, if ligand word 1 activates cell type A, while ligand word 2 activates cell types B and C, then (A) and (B, C) constitute distinct channels.

### Combinatorial addressing(or simply addressing)

A mapping between ligand words and the corresponding cell type(s) activated by those words.

### Orthogonal addressing

A particular form of combinatorial addressing in which each ligand word activates a single, unique cell type. An example of 3-channel orthogonal addressing is shown in Figure 3B.

### Addressing repertoire

The combinations of cell types (each combination representing a channel) that can be activated across all possible ligand words for a given set of cell types and biochemical parameters. Examples of addressing repertoires are shown with the Venn diagrams in Figure 5A.

### Bandwidth

The number of unique channels in a given system. For example, a system where ligand words 1 and 2 both activate only cell type A while ligand word 3 activates cell type B would have a bandwidth of 2, as ligand words 1 and 2 yield the same activation profile.

The next two terms define quantitative metrics used in this paper.

### Distinguishability

(Figures 3–5, Methods: Distinguishability of Orthogonal Channels) quantifies the specificity of a given addressing scheme and is defined as (lowest on-target)/(highest off-target) activity. For example, suppose ligand word 1 activates on-target cell type A and off-target cell type B at levels of (0.8, 0.1) units, respectively, while ligand word 2 activates off-target cell type A and on-target cell type B at levels of (0.4, 0.9). The distinguishability for orthogonal addressing of (A) and (B) is 0.8/0.4 = 2. As another example, if cell types A and B are both on-target for ligand word 2, addressing (A) and (A, B) would have a distinguishability of 0.4/0.1 = 4.

### Addressability

(Figure 6, Methods: Addressability Metric of Ligand Words) quantifies the diversity of the addressing repertoire for a set of ligand words. We first measure the separation of two ligand words as the largest ratio of their resulting activation levels in any cell type. Addressability is then defined as the separation of the least separable pair of ligand words.

## Supplemental Figures

**Figure S1:**
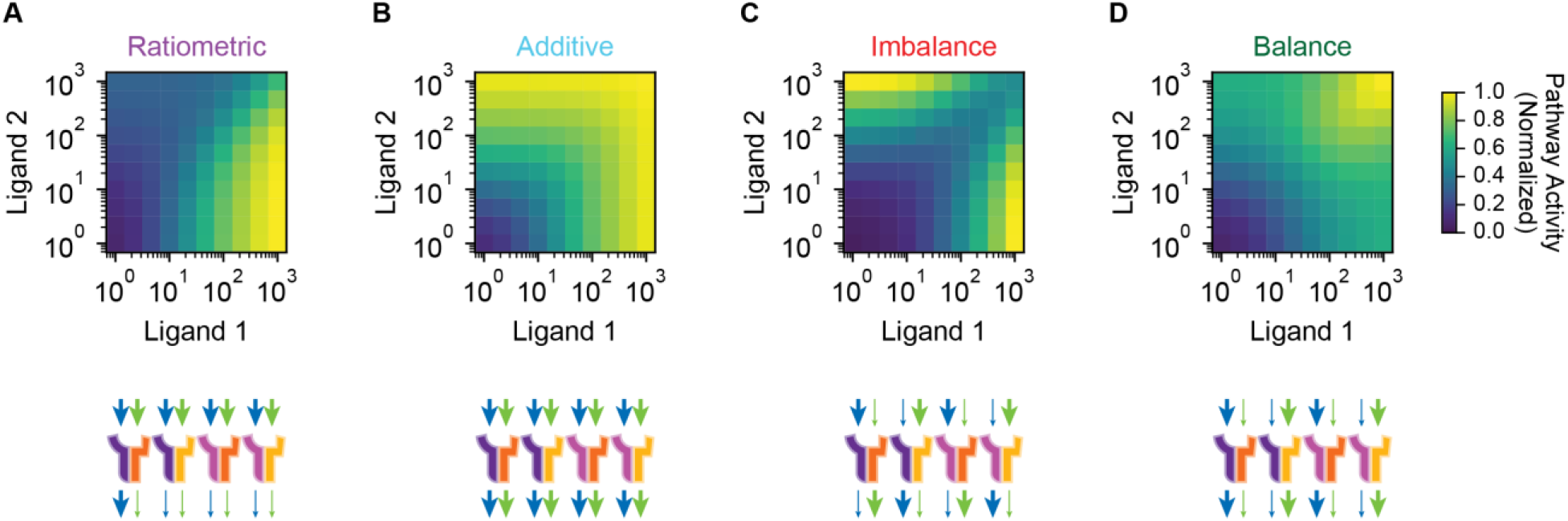
Promiscuous ligand-receptor interactions generate a repertoire of archetypal response functions. Four archetypal response functions — ratiometric, additive, imbalance, and balance — appeared in a more complex model of the BMP signaling pathway (Antebi et al., 2017). Here, we show that similar archetypes appear in the model used here (top), along with the model parameters that generate them (bottom). Parameter diagrams represent the affinity (top arrows) and activity (bottom arrows) parameters corresponding to each signaling complex. Arrow width indicates relative magnitude. Thin arrows correspond to values of 0.1, while thick arrows represent values of 1. (A) In ratiometric responses, one ligand reduces the activity of the other, such that the overall response approximates the ratio of the two concentrations. Such responses can arise through competitive inhibition, where a second ligand binds the receptors that are needed to generate signaling activity but produces inactive signaling complexes. (B) Additive responses approximate the sum of the two ligand concentrations, as the ligands increase pathway activity either alone or together. Ligands that activate receptors equivalently can generate such responses. (C) In imbalance detection, the pathway is most active when there is a large imbalance in the levels of the two ligands. These responses can arise if, for instance, competition between two ligands favors complexes with low signaling activity. (D) Balance detection responses show most activity when both ligands are present simultaneously at a particular ratio. One mode for generating them is when ligand binding favors formation of high-activity signaling complexes.

**Figure S2:**
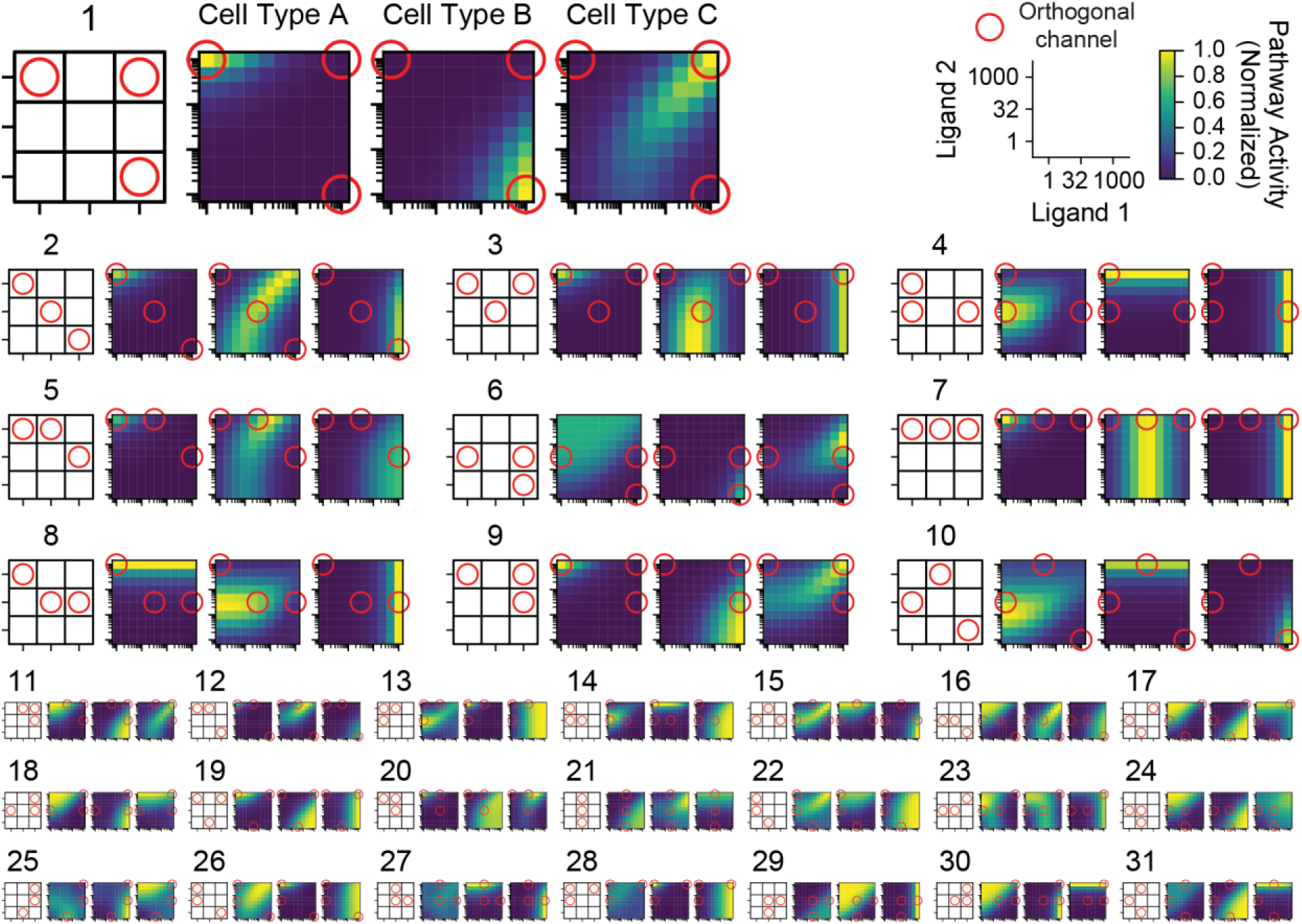
Orthogonal addressing can arise from a variety of different response types. For the parameter sets represented in Figure 3D, the responses of each cell type are shown. Parameter sets are ordered by distinguishability, from best to worst. These responses illustrate that three-channel addressing can be achieved in a variety of ways, although common patterns do emerge (cf. schemes 1-2 and schemes 3-4). Different scales are used to focus on the strongest examples.

**Figure S3:**
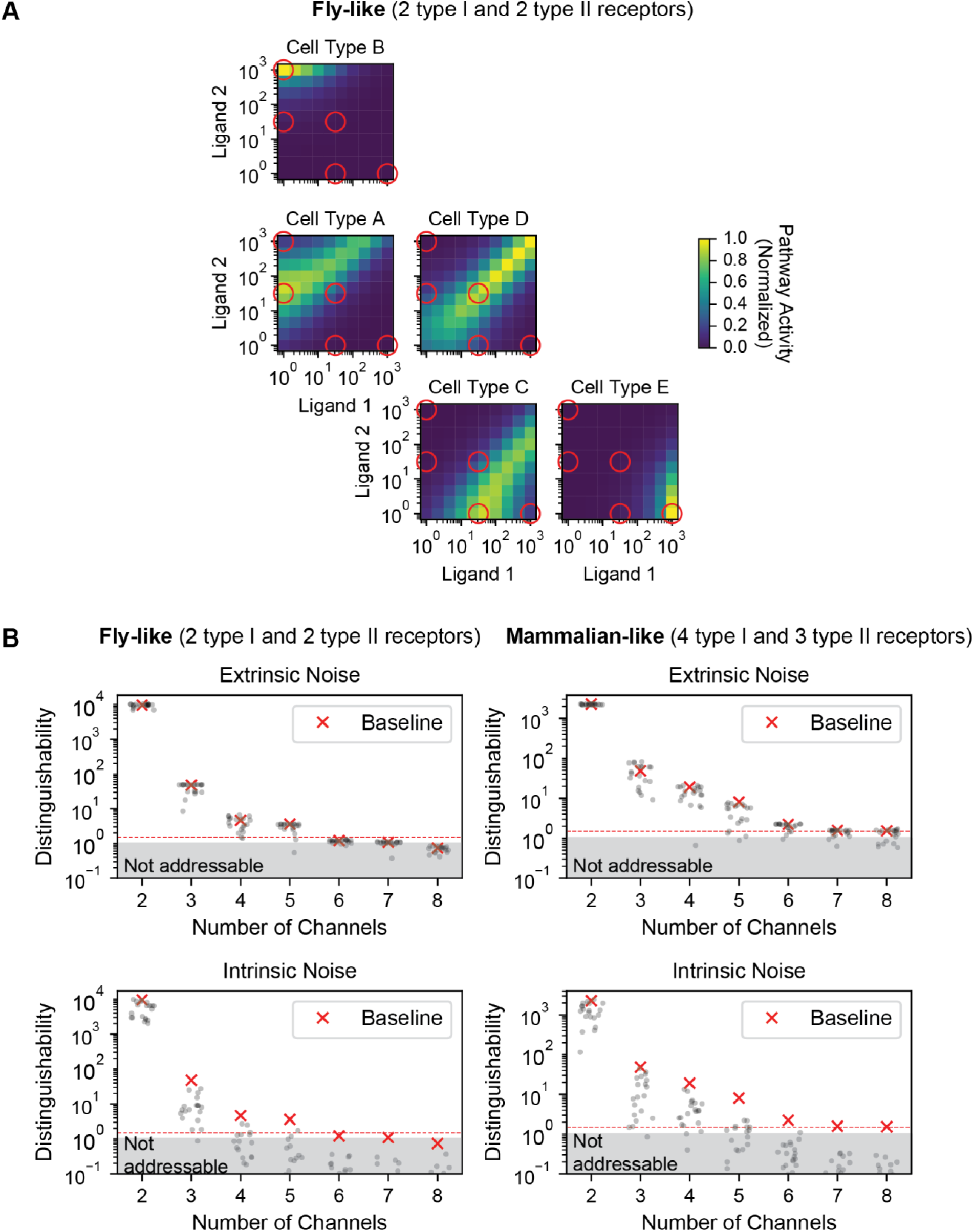
Orthogonal addressing schemes are robust to extrinsic noise in receptor expression levels. (A) The responses of each cell type in the 5-channel system analyzed in Figures 4C-D are shown. As in Figure 4I, ligand words corresponding to orthogonally activating channels are shown as red circles. Responses have been rearranged such that the response of a given cell type is shown in the relative position of its orthogonally activating ligand word. For example, the top left cell type is activated by high levels of ligand 2 only, while the bottom right cell type is orthogonally activated by high levels of ligand 1 only. (B) Top parameter sets from the fly-like model (2 type I and 2 type II receptor variants) of Figure S3A (left) and the mammalian-like system (4 type I and 3 type II receptor variants) of Figure 4I (right) were evaluated for addressability in the presence of noise. Receptor expression levels were perturbed with extrinsic (correlated, top) or intrinsic (uncorrelated, bottom) noise (Methods: Analysis of Robustness), and distinguishability values were computed with all other parameters held constant. For each condition, the results of 20 perturbations are shown, along with the baseline value (red crosses).

**Figure S4:**
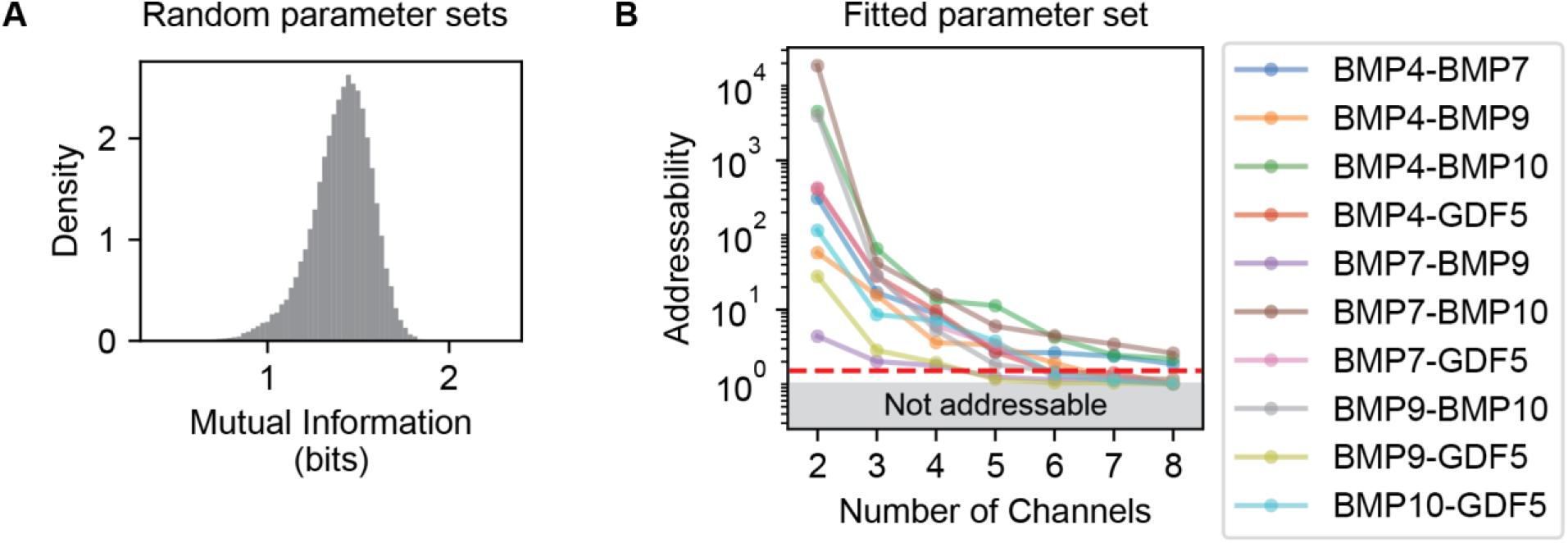
Addressing properties vary across parameter sets. (A) The approach of Figures 6A-B was applied to an equivalent number of randomly generated parameter sets rather than a grid of parameter values. The resulting distribution of mutual information values is similar, indicating that the result is robust to the method of parameter sampling (cf. Figure 6B). (B) Parameters for 5 ligands, 2 type I, and 3 type II receptors were fitted to experimental measurements of BMP responses in multiple cell lines with differing receptor expression profiles (Klumpe et al., 2020) and analyzed for their addressing potential. Specifically, we computed the addressability for every pair of ligands using the same libraries of ligand combinations and cell types as in (A). Different pairs show varying levels of addressing potential, with some pairs (BMP4-BMP7, BMP4-BMP10, and BMP7-BMP10) exhibiting addressing of different cell type groups for every ligand combination and others (e.g. BMP7-BMP9) showing lower bandwidths.

## Methods

### Resource Availability

Further information and requests for resources and reagents should be directed to and will be fulfilled by Michael B. Elowitz. The data and code generated during this study are available at http://dx.doi.org/10.22002/D1.1692.

### Experimental Model and Subject Details

#### Cell Lines

NAMRU mouse mammary gland (NMuMG) cells (female) were acquired from ATCC (CRL-1636). Mouse embryonic stem cells (mESCs; E14Tg2a.4) were obtained from the laboratory of Bill Skarnes and Peri Tate. Reporter cell lines were generated and cultured as in (Antebi et al., 2017). Briefly, cells were cultured in a humidity-controlled chamber at 37^*◦*^C with 5% CO_2_. NMuMG cells were cultured in DMEM supplemented with 10% FBS (Clontech #631367), 1 mM sodium pyruvate, 1 unit/ml penicillin, 1 *μ*g/ml streptomycin, 2 mM L-glutamine, and 1× MEM nonessential amino acids. mESCs were plated on tissue culture plates pre-coated with 0.1% gelatin and cultured using DMEM supplemented with 15% FBS (Gibco #16141), 1 mM sodium pyruvate, 1 unit/ml penicillin, 1 *μ*g/ml streptomycin, 2 mM L-glutamine, 1× MEM nonessential amino acids, 55 mM *β*-mercaptoethanol, and 1000 units/ml leukemia inhibitory factor (LIF).

### Method Details

#### Addressing of Cell Lines

##### siRNA-Induced Knockdown

Cells were plated at 40% confluency in a single well of a 24-well plate with 30 *μ*M total siRNA (ThermoFisher *Silencer* Select #4390771) and 3 *μ*l RNAiMAX (Invitrogen). For every gene, we used a pool of two distinct siRNAs (*Acvr1*, Lifetech #S61924 and #S61925; *Bmpr2*, Lifetech #S63047 and #S63048). Cells were passaged after 24 hours and were used for the specified experiments.

##### BMP Response and Flow Cytometry

Cells were plated at 40% confluency in 96-well plates and cultured under standard conditions for 12 hours. Media was then replaced and ligands were added at specified concentrations. 24 hours after ligand addition, cells were prepared for flow cytometry by washing with PBS and lifting from the plate using either trypsin (NMuMG) or Accutase (mESC) for 5 minutes at 37°C. Protease activity was quenched by resuspending the cells in HBSS with 2.5 mg/ml bovine serum albumin (BSA). Cells were then filtered with a 40 mm mesh and analyzed by flow cytometry (MACSQuant VYB, Miltenyi). Recombinant BMP ligands were acquired from R&D Systems (BMP4, #5020-BP; BMP9, #5566-BP).

Single-cell flow cytometry data were analyzed as in (Antebi et al., 2017) by taking the population median. For measured experimental responses (Figure 2), responses were measured by taking the mean of at least 3 repeats. Relative activity represents fold change compared to response at lowest concentrations of each ligand.

#### One-Step Model for Promsicuous Interactions

##### Ligand-Receptor Interactions

Many signaling pathways demonstrate promiscuous interactions between multiple ligand and receptor variants, which can bind with varying affinities to form many distinct signaling complexes. The BMP pathway represents a canonical example of such an architecture. Previously, we have described a mathematical model that captures key features of this pathway and recapitulates experimentally observed responses (Antebi et al., 2017). Here, we develop a simplified version of the model that captures equivalent behaviors at steady state while reducing the number of parameters to be considered.

In the model, we describe binding of a ligand to a heterodimer of type I and type II receptors. Specifically, we consider *n_L_* ligand variants, *n_A_* type I or A receptor variants, and *n_B_* type II or B receptor variants, where ligand *L_i_* can interact with A receptor *A_j_* and B receptor *B_k_* to form the heterotrimeric signaling complex *T_ijk_*. We assume that this process occurs as a one-step reaction with an effective three-way interaction, with forward rate 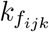 and reverse rate 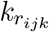. This reaction can be summarized as follows:

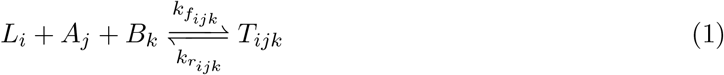

##### Differential Equations and Constraints

Letting *L_i_* denote the concentration of ligand in a volume *V* and *A_j_*, *B_k_*, and *T_ijk_* denote the absolute numbers of receptors and complexes on the cell surface, we can then write the differential equations that describe the dynamics of these reactions:

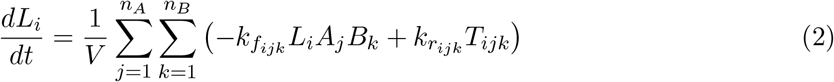

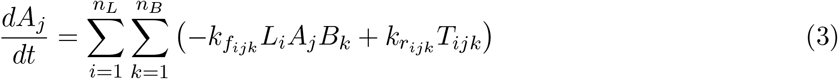

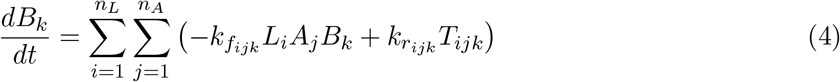

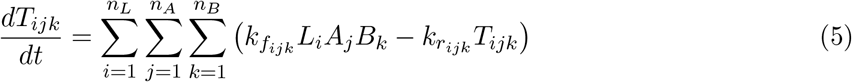

Each complex *T_ijk_* phosphorylates the second messenger at some rate *∊_ijk_* to generate intracellular signal *S*, which degrades at rate *γ*. The rate of change of the total signal is given by the following differential equation:

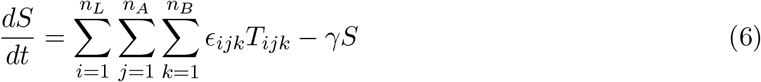

We assume that the volume for the ligands is large, or *V* → *∞*. In this regime, there are significantly more ligand molecules than receptors, as is the case for experimental conditions in which ligands are dissolved in an excess of media. Under this assumption, ligand concentrations remain constant. We further assume that production and consumption of the various molecular species are in steady state. By conservation of mass, the total number of each type of molecule (alone or in complex with other species) must remain constant. Letting 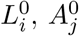, and 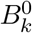 denote the initial values of the respective species, we obtain the following constraints:

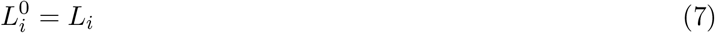

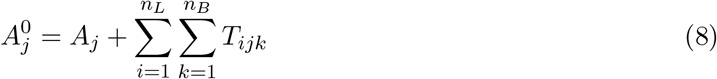

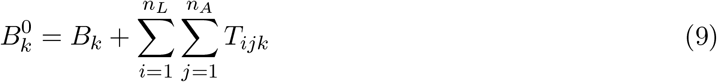

##### Steady-State Equations

Since binding and unbinding of ligands and receptors occur on fast time scales relative to the time scales of reporter detection, we focus on characterizing the behavior of this system at steady state. Here, all time derivatives in eqs. (2) to (6) vanish. Defining affinities 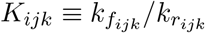 and activities *e_ijk_ ≡ ∊_ijk_/γ*, eqs. (5) and (6) can be solved as follows:

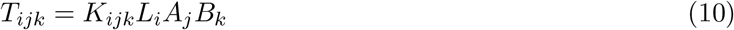

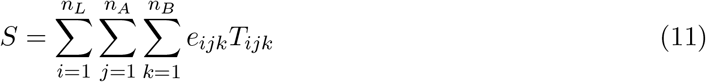

Together, eqs. (7) to (11) describe the behavior of the model at steady state.

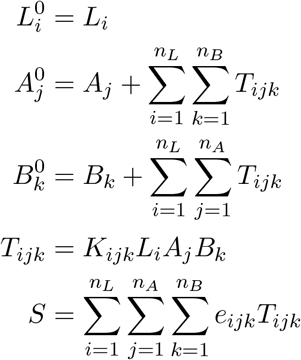

We can solve this system of equations to find the values of *T_ijk_* at steady state, which we can then use to compute the total signal *S*. From eqs. (8) and (9), the steady-state values of the receptors are as follows:

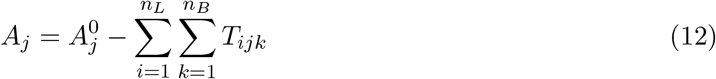

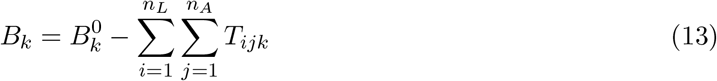

Substituting into eq. (10), we have a system of *n_T_* = *n_L_n_A_n_B_* quadratic equations for *T_ijk_*:

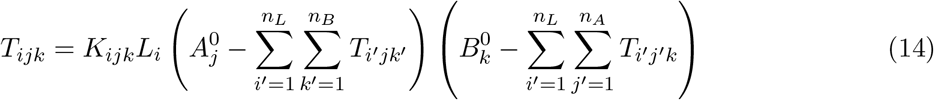

The solutions for *T_ijk_* from this system of equations can then be substituted into eq. (11) to compute the total signal *S*.

To solve the model efficiently, we used Equilibrium Toolkit (EQTK) (Bois, 2020), an optimized Python-based numerical solver for biochemical reaction systems. EQTK casts the coupled equilibrium problem as an unconstrained convex dual optimization problem and employs a globally convergent trust region algorithm to solve it (Bois, 2020; Dirks et al., 2007). This method accelerated computation by 600-fold compared to standard nonlinear least-squares optimization used previously (Antebi et al., 2017).

#### Comparison with Alternative Models

##### One-Step vs. Two-Step Model

We have previously considered a mathematical model that describes the promiscuous architecture of the BMP pathway, which considers formation of the heterotrimeric complexes *T_ijk_* in a two-step process (Antebi et al., 2017). Briefly, ligand *L_i_* and receptor *A_j_* form an intermediate dimer *D_ij_*, which then binds to receptor *B_k_* to form trimer *T_ijk_*. Again, we assume that the reactions are reversible and follow first-order kinetics, with forward and reverse reaction rates 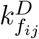 and 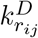 for formation of the dimers and 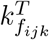 and 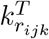 for formation of trimmers. Defining 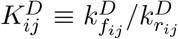 and 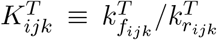, the steady-state solutions for *T_ijk_* in the two-step model, analogous to, eq. (14) in the one-step model, are as follows:

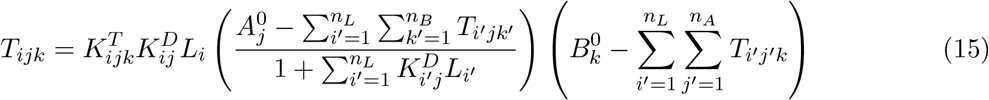

Comparing eq. (14) and eq. (15), the steady-state solutions for *T_ijk_* in the two-step model can be mapped to the one-step model under the following parameter choice:

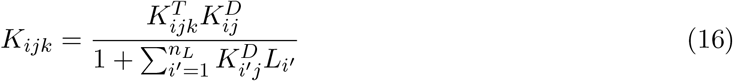

Since *S* is defined by the values of *T_ijk_* and is given by eq. (11) in both the one-step and two-step models, the steady-state behavior of the two-step model with any set of parameters can also be represented in the one-step model. However, the number of parameters is reduced from 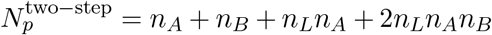 to 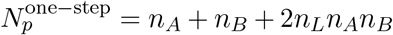. Thus, the one-step model enables us to simplify the system while preserving all possible behaviors of *T_ijk_* and *S* at steady state.

##### Trimeric vs. Hexameric Model

We have developed a simplified model in which a ligand binds to type I and type II receptor subunits to form a trimeric signaling complex. However, the BMP signaling pathway is known to involve hexameric signaling complexes, where a dimeric ligand interacts with two type I and two type II receptors. This model captures reactions of the following form:

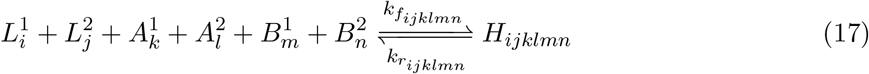

This model can essentially be reduced to a trimeric model by setting reaction rates to 0 for any reaction with *i* ≠ *j*, *k* ≠ *l*, or *m* ≠ *n*. As such, responses in the trimeric model represent a subset of the functions that could be possible in the hexameric model.

#### Optimization of Orthogonal Addressing Schemes

Given a target orthogonal addressing scheme of *N* channels, we optimized for parameters that would yield matching responses. Specifically, we used constrained least-squares optimization for *n_L_n_A_n_B_* affinity parameters *K_ijk_*, *n_L_n_A_n_B_* activity parameters *e_ijk_*, and *N* (*n_A_* + *n_B_*) receptor expression levels. We bounded affinity and efficiency parameters in [0, 1] and receptor levels in [0*, ∞*). We sought to minimize the residuals between the target responses and the simulated responses at the *N* ligand words of interest. Since the simulated responses have arbitrary units, we normalized all responses for a given parameter set. Specifically, we normalized by the maximum value in any cell type over the full ligand titration, not only the ligand words of interest. This normalization ensures that all cell types share a relatively similar level of activation and that the activation in the orthogonal channels is distinguishable from activation by other ligand combinations.

As this optimization procedure is not guaranteed to converge to a global minimum, we optimized repeatedly with different initial conditions. All parameters were chosen in a uniform random distribution over [0, 1].

#### Distinguishability of Orthogonal Channels

We optimized parameters for each addressing scheme based on the squared error between the targeted and simulated responses at each ligand word of interest. However, this error does not necessarily guarantee specificity of addressing, where a given ligand combination should activate only a single cell type and not the others. To quantify the performance of each system, we visualized the distributions of on-target and off-target activation levels. We defined the distinguishability as the fold difference between the minimum on-target and maximum off-target activities, which measures the ability to differentiate between specific and nonspecific signals in the worst case.

#### Enumeration of Orthogonal Addressing Schemes

To analyze the possibility for orthogonal addressing, we used a discretized ligand concentration space to enable a comprehensive screening. We reasoned that we could systematically test for all possible orthogonal addressing schemes by selecting a subset of the possible ligand combinations to be orthogonally activating and defining a set of targeted response functions accordingly. For a set of *N* chosen ligand words, we enumerated *N* targeted response functions, where each ligand word activates exactly one cell type and, conversely, each cell type is activated by exactly one ligand word. Having discretized ligand concentrations to 3 levels, there were 3^2^ = 9 possible ligand words. Since the combination with all ligands at the lowest level is assumed to yield negligible activation in any cell type, there can be 1 to 8 possible communication channels. Therefore, for each possible number of channels *N*, we took all possible subsets and sought to achieve these addressing schemes. There are 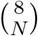 possible addressing schemes for a given bandwidth and 2^8^ = 256 possible addressing schemes overall. However, addressing schemes that are identical under changes in ligand labels were removed, to give 144 total possibilities.

To evaluate the potential capacity of promiscuous ligand-receptor systems for orthogonal addressing, we sought to optimize progressively higher bandwidths. For *N* channels, we randomly selected *N* ligand words as orthogonally activating inputs and sought to optimize parameters as described above. We iterated this process with randomly chosen ligand words until parameters had been identified generating distinguishability of at least 2. Once this criterion was met, we then proceeded to optimize *N* + 1 channels, up to the limit of 8. We performed 10, 000 optimizations.

To more systematically test for the ability to achieve each addressing scheme, we also performed a search over all schemes rather than considering only the resulting bandwidth. Specifically, we chose which addressing scheme to optimize based on both the lowest error *E* achieved and the number of trials *n* attempted. Specifically, we optimized for the addressing scheme with maximal value of *E/n*^2^ and repeated this process over 7, 200 trials (an average of 50 optimizations per scheme) to ensure that all schemes would be tested adequately.

#### Analysis of Robustness

Biological systems are subject to noise. In particular, cellular systems show both extrinsic noise, or correlated changes such as during cell growth or changes in expression machinery, and intrinsic noise, or independent stochastic variation in each element. To assess whether the optimized parameters are robust to noise in receptor expression levels, we evaluated whether on-target and off-target signals could be correctly distinguished across many random perturbations, using the receiver operating characteristic (ROC). Specifically, we computed the area under the ROC curve (AUC), which represents the probability of successfully classifying on-target from off-target activations. We considered both purely extrinsic and purely intrinsic noise. For a given coefficient of variation (CV) *ν* (here, *ν* = 0.5), we simulated extrinsic noise by generating a scale factor *s* from a gamma distribution with shape parameter 1*/ν*^2^ and scale parameter *ν*^2^ (giving a mean of 1 and a variance of *ν*^2^) and multiplied all receptor levels by this scale factor. For intrinsic noise, we instead drew scale factors i.i.d. for each receptor. With each form of noise, we generated 100 random perturbations, simulated the resulting activity levels, and computed the corresponding ROC and AUC (Figures 4D,H). We also plotted the distinguishability values for 20 such perturbations (Figure S3B).

#### Addressing Repertoires

##### Enumeration of Addressing Repertoires

We next considered more general addressing systems, targeting not just activation of individual cell types but also groups of cell types. Specifically, an addressing repertoire encompasses all subsets of cell types that can be coactivated by a ligand word, across a complete titration of ligand concentrations. For instance, titrating two ligand variants with three concentrations yields nine ligand words, each of which activates some subset of the cell types considered. Every distinct group of cell types constitutes an achievable channel. The set of channels resulting from the nine ligand words considered constitutes the addressing repertoire for that set of parameters.

We focused on analyzing addressing repertoires for three cell types, denoted A, B, and C. With three cell types, there are eight possible channels: one with no cell types activated, three with a single cell type activated, three with two cell types activated, and one with all cell types activated. Since the channel with no cell types activated is always achieved in the absence of any ligand, we neglect this from further consideration. Each of the remaining seven channels may or may not be present, for a total of up to 2^7^ = 128 addressing repertoires. We discard repertoires that are invariant with respect to relabeling of ligands as well as repertoires in which two cell types are indistinguishable by any ligand combination (such that the addressing repertoire could be mapped to one for two cell types). As discussed in more detail below, these simplifications leave us with 32 addressing repertoires of three cell types, corresponding to those shown in Figure 5C.

We first seek to eliminate repertoires that are invariant with respect to relabeling of ligands. The subset with all three cell types activated, or “triple,” does not change when ligands are relabeled; however, singles or doubles may. For example, the addressing repertoire consisting of “A” and “BC” is equivalent to that comprising words “B” and “AC,” simply by swapping cell types A and B. As such, we consider the unique ways to include singles or doubles. Consider each single with its complementary double (for example, “A” with “BC”). There are 2^2^ = 4 possible ways to include this pair in an addressing mode: both absent, only single present, only double present, and both present. The three pairs can then have three distinct choices (4 combinations), two distinct choices (4 · 3 = 12 combinations), or the same choice (4 combinations). There are then 4 + 12 + 4 = 20 ways to choose combinations of singles or doubles, and the triple may be either present or absent. Thus, considering ligand relabeling reduces the total number of addressing repertoires to consider to 40.

Note, however, that some of these repertoires may only represent two distinct cell types, rather than three. For example, the addressing repertoire with channels “A” and “BC” indicates that cell types B and C are indistinguishable across all ligand combinations and are therefore functionally equivalent. Let B and C be indistinguishable, without loss of generality. The only possible channels are then “A,” “BC,” and “ABC.” Thus, the 2^3^ = 8 repertoires that only contain these channels can be reduced to two distinct cell types and are therefore omitted from our analysis of addressing three cell types. This correction yields our final set of 32 addressing repertoires.

Similar to a specific number of orthogonal channels, a given addressing repertoire can potentially be implemented in many ways. In other words, many different sets of responses can generate the same addressing repertoire. Unlike the orthogonal case, however, the responses for every ligand combinations are relevant. Thus, enumerating the ways to achieve a given addressing repertoire requires considering any possible response for every cell type.

##### Optimization of Addressing Repertoires

To generalize our optimization approach to analyze addressing repertoires, we first set out to define what sets of responses could yield a given repertoire. Therefore, we started by enumerating all possible binary response matrices for a single cell type. The number of possible responses then reduces to the number of ways to choose “on” signals. Assuming that cells are always inactive for the ligand combination where both ligands are present at low levels and ignoring the case where the cell is entirely nonresponsive, there are 2^8^ − 1 = 255 possible responses.

By considering all combinations of three responses from this set, we were able to map all addressing repertoires to the potential sets of three responses. Due to the large number of possibilities for a given repertoire, we sought to prioritize sets of responses that were more likely to be achievable. Therefore, we individually optimized each of the 255 responses and quantified the quality of each response using the sum of squared distance to the target, after normalizing the simulated response to have a maximum response of 1 (data not shown). We ranked sets of three responses based on the sum of the scores of each response individually. Since parameter sets were individually optimized, a response that can be achieved with high quality independently may not be possible in the same biochemical parameter regime as another; however, this scoring should reduce consideration of responses that are challenging to optimize individually, let alone together with others.

Having selected candidate sets of responses, we could then perform least-squares optimization as done previously. We also complemented this optimization approach by reasoning that any given set of responses matches some addressing repertoire, depending only on how the threshold between “inactive” and “active” pathway response is defined. Therefore, we simulated a random set of responses, chose the threshold that yielded the greatest distinguishability between the lowest on-target and highest off-target responses, and associated those parameters with the resulting addressing repertoire. We iteratively optimized for this distinguishability, stopping if the resulting addressing repertoire was one for which a valid parameter set had not yet been identified.

To characterize the specificity of addressing different subsets of cell types, we generalized the distinguishability metric defined above. Each ligand word activates a particular subset of cell types; the corresponding response(s) of the cell type(s) would represent on-target signaling, while the response(s) of any other cell type(s) would represent off-target signaling. Therefore, as in the case of orthogonal addressing, distinguishability can be calculated as the fold difference between the minimum on-target activity and the maximum off-target activity.

##### Comparison with One-to-One Architecture

To understand how subset addressability in a promiscuous pathway compares with that in a one-to-one architecture, note that all responses in a one-to-one pathway must be monotonic, meaning that responses never decrease with added ligand. As such, a given cell’s response is maximal when exposed to the ligand combination where all ligands are present at highest concentration. Therefore, every cell type will be active in response to this ligand combination. (Otherwise, there would be no response across the entire ligand titration, and there would be no “addressing.”) Consequently, the subset “ABC” will always be addressable in the one-to-one architecture. Conversely, any repertoire where “ABC” is absent cannot be achieved in the one-to-one architecture.

Additionally, a one-to-one architecture can only generate as many orthogonal channels as there are ligand variants, due to the monotonicity of responses. Therefore, any addressing repertoire that enables orthogonal addressing of three cell types cannot be achieved with only two ligands.

#### Computation of Mutual Information

##### Mathematical Framework

We use mutual information between ligand words and activation patterns across a library of cell types to quantify the combinatorial addressing power of the ligand-receptor system. Mutual information was initially developed to quantify the capacity of a noisy channel to transmit information, or the extent to which distinct input messages can be resolved by the receiver after passing through the channel. Here, we view the ligand words as input messages and the resulting activation pattern across cell types as the received message. Then, the communication system’s capacity is determined by the biochemical constants *K_ijk_* and *e_ijk_*.

One significant benefit of using an information theoretic framework is that we do not need to assume a particular set of ligand words and cell types and then optimize over them. Instead, we can use extensive libraries of input ligand words and cell types; mutual information will reflect the best subset of each with no penalty (or benefit) for redundancies. Thus, in our framework, mutual information reflects a property of the biochemical constants *K* and *e* alone; ligand inputs and cell types are implicitly assumed to be optimally chosen. (In information theoretic language, we do not need to know optimal error-correcting codes to compute the capacity of a channel.)

Let **c** represent a library of *n_LW_* ligand words, where the *i*th input **c**_*i*_ is a vector of *n_L_* ligand concentrations. Given a library of *n_CT_* cell types, let **a**(**c**) represent the resulting activation profiles of these cell types, or a set of *n_LW_ × n_CT_* responses. Earlier sections have presented a way to compute **a**_determ_(**c**) deterministically by solving quadratic equations. Here, we assume that the activation is probabilistic due to a Gaussian error bar of size *σ* around **a**_determ_(**c**); the standard deviation *σ* can represent molecular fluctuations upstream of SMAD (e.g., in receptor levels or activity) that result in fluctuations of SMAD phosphorylation. Thus,

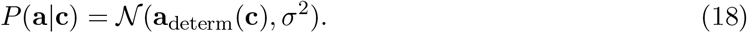

We compute mutual information MI(**a**, **c**) using the formula

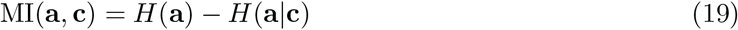

The second term can be expressed as

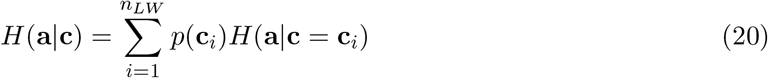

Each *H* (**a***|***c**= **c**_*i*_) is the entropy of a *n_CT_*-dimensional Gaussian with covariance matrix Σ = *σ*^2^*I*, where *I* is the identity matrix of size *n_CT_*. This entropy (in bits) is

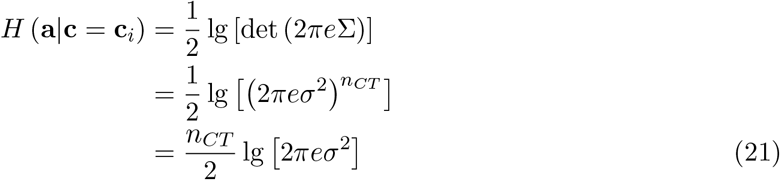

Assuming input probabilities are uniformly distributed, or 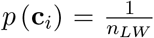, this conditional entropy is simply

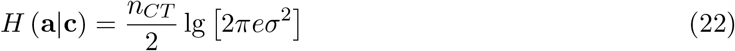

Thus, this term is constant regardless of choice of biochemical parameters.

The entropy *H*(**a**) in eq. (19) is the entropy of *P* (**a**), which is a sum of Gaussians, one at each of the activation patterns corresponding to each ligand input **c**_*i*_. This entropy is a measure of the distinguishability of activation patterns **a**(**c**_*i*_) for different inputs **c**_*i*_; the entropy will be small if the Gaussians are overlapping and large otherwise. Intuitively, this entropy is a measure of how well-separated the activation patterns **a** for different ligand inputs are.

The problem of determining the entropy of a normalized sum of Gaussians (i.e., a Gaussian mixture) in high dimensions is surprisingly involved; however, simple analyic approximations have been developed in a recent advance (Kolchinsky and Tracey, 2017). We use the approximation to the kernel density estimator presented therein for a sum of *n* Gaussians 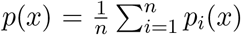 in *d* dimensions:

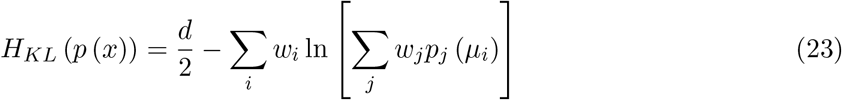

Here, *p_j_* is the *j*th Gaussian component (normalized to 1, individually), *μ_i_* the mean of the *i*th component, and *w_i_* the mixture weight of the *i*th component. In this case, we assume uniform mixture weights, or 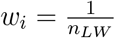 for all *i*. Further, *p_j_* (*μ_i_*) is

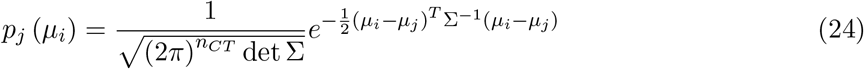

Thus, the mutual information can be evaluated by simply evaluating each Gaussian at the mean of all other Gaussian components, or using the matrix *D_ij_* of distances between activation patterns **a**_determ_ (**c**_*i*_) for different ligand inputs **c**_*i*_:

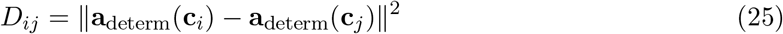

We can therefore simplify *p_j_* (*μ_i_*) to

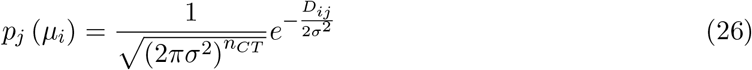

Substituting into the approximation, we find an entropy (in nats) of

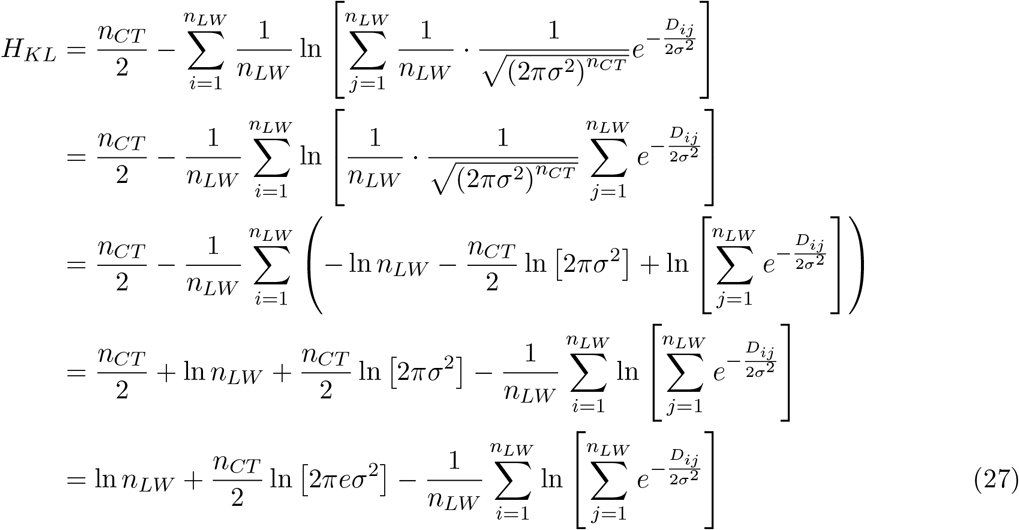

We can convert this expression to bits by multiplying by lg 2 and combine with eq. (22) to estimate mutual information. We note that these derivations omit a correction factor *C* = *−n_CT_* lg Δ*a* arising from binning with bin width Δ*a* to make the entropy of a continuous distribution well defined; however, as the same correction applies to both *H* (**a**) and *H* (**a**|**c**), these terms cancel out.

Our estimator of mutual information is therefore

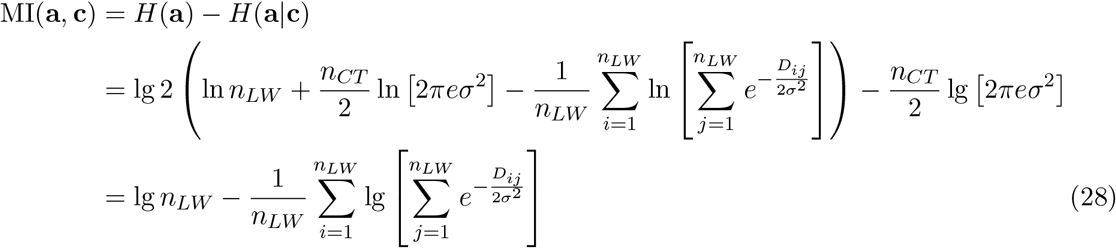

We use this expression to estimate mutual information in this paper. From the form, it is clear that mutual information rewards large values of *D_ij_*, i.e., distinct activation patterns for different ligand inputs.

The mutual information framework above can be naturally extended to scenarios not considered here. For example, not all ligand inputs might be equally likely or of equal physiological significance. In this case, the map of ligand inputs to activation profiles (i.e., the coding scheme) can separate the activation patterns of more important ligand words at the expense of more similar activation patterns for less important words. The mutual information framework can account for such weighting of different inputs easily through unequal *p*(**c**_*i*_) above.

Finally, note that mutual information naturally rewards robustness, since mutual information is higher when each activation pattern is realized over equally sized regions of input space. For example, if an output **a**_1_ is only obtained for a sliver of ligand input space while another output pattern **a**_2_ is realized over the rest of input space, mutual information will be lower than if both outputs are realized over half of input space.

#### Libraries of Ligand Words and Cell Types

To provide the broadest information-theoretic characterization, we first constructed comprehensive libraries of ligand words, or input messages, and cell types, or receptor expression profiles. Each ligand can independently take on three distinct concentrations sampled logarithmically over three orders of magnitude, or 10^0^ = 1, 10^1.5^ *≈* 32, and 10^3^ = 1000. This library of 3^2^ = 9 words is a representative sampling of all possible ligand inputs. Similarly, we constructed a library of cell types by varying each receptor level independently over two distinct concentration levels [0.1, 1]. For a system with 2 type I receptor variants and 2 type II receptor variants, this library comprises 2^2+2^ = 16 possible cell types.

#### Sampling of Biochemical Parameters

In our model, a promiscuous ligand-receptor system is defined by its interaction affinities *K* and signaling activities *e*. To comprehensively sample all possible biochemical parameters, each parameter was allowed to be either of [0.1, 1], giving a total of 2^16^ = 65,536 qualitatively distinct parameter sets. We then evaluated mutual information between the ligand words and the corresponding cell type activation profiles for each possible choice of biochemical parameters *K, e*. The resulting data characterize the combinatorial addressing power across a comprehensive set of promiscuous ligand-receptor systems.

##### Random Sampling of Biochemical Parameters

To ensure that the lattice sampling procedure did not introduce any artifacts, we also repeated this analysis for an identical number of randomly generated parameter sets. Specifically, we chose each value independently and randomly with a log-uniform distribution over [10^−1.5^, 1]. The resulting distribution of mutual information values is shown in Figure S4A.

##### Choice of Variance

Computing mutual information requires choosing the variance or Gaussian fluctuation *σ* for all activation levels. For all results here, we choose *σ*^2^ = 0.5 after testing a range of values. Very large or small choices of *σ* lead to the same value of mutual information for all biochemical parameters, either low or high, respectively. Intermediate choices of *σ* discriminate between different *K, e*. While the precise value of mutual information depends on *σ*, different choices of *σ* do not qualitatively change the relative ordering of biochemical parameter sets.

##### Optimization of Mutual Information

These sampling procedures enable us to comprehensively analyze mutual information across parameter space. However, they are likely to miss extremes of mutual information. Therefore, we chose 16 parameter sets from the lattice sampling with the highest starting mutual information and further refined *K, e* to maximize mutual information using least-squares optimization.

#### Addressability Metric of Ligand Words

To determine which ligand words in the library activate distinct combinations of cell types in this optimal limit, we define the overall addressability of all ligand words by evaluating all pairs. To compare a pair of ligand words, we compute the ratio of activation levels for each cell type and take the separation *r* as the largest such fold change (inverted if needed, such that *r* ≥ 1). If two ligand words have a separation of 5, then at least one cell type’s activation is different by a factor of at least 5 in the two conditions. We extend this pairwise separation to a set of *N* ligand words by forming a *N* × *N* addressability matrix, where element (*i, j*) corresponds to the pairwise separation of ligand words *i* and *j*. This matrix has 1s along the diagonal. We define the smallest off-diagonal value, which represents the minimum pairwise separation between different ligand words, to be the overall addressability of that set of *N* ligand words.

#### Analysis of Archetypal Responses

##### Response Classes

We next analyzed the responses generated by the full library of cell types for high-performing and low-performing parameter sets. As expected, parameter sets giving rise to low mutual information showed relatively little diversity in responses (Figure 6D). Parameter sets which generated high mutual information showed distinct activation patterns among cell types (Figure 6E); furthermore, these response types appeared qualitatively different and were similar to experimentally observed patterns reported previously (Antebi et al., 2017). We therefore further analyzed the presence of these archetypal responses across parameter sets.

Examples of these archetypes were generated by simulating responses to parameters reflecting our understanding of the underlying design principles (Figure S1). These parameters are not specifically tuned, with all affinity and activity values set to either 0.1 or 1 and all receptor levels fixed at 10^*−*1.5^. Thus, they reflect qualitative differences rather than finely tuned quantitative ones. Briefly, ratiometric responses feature reduction of activity of one ligand by the second, such that the overall response approximates the ratio of the two concentrations. Competitive inhibition, where the “denominator” competes for receptors needed to generate signaling activity but produces inactive complexes, can produce such responses (Figure S1A). Additive responses approximate the sum of the two ligand concentrations, as the ligands increase pathway activity either alone or together, and are readily generated when both ligands activate receptors similarly (Figure S1B). Imbalance detection responses, where cells respond maximally to imbalances in the levels of the two ligands, can arise if, for instance, competition between two ligands favors complexes with low signaling activity (Figure S1C). Conversely, balance detection responses, where cells respond maximally when both ligands are present at a specific ratio, can be generated when ligand binding favors formation of high-activity signaling complexes (Figure S1D).

##### Phenotypical Parameters

We characterized the spectrum of responses as described previously (Antebi et al., 2017). Briefly, we use the relative ligand strength (RLS), which represents the ratio of activation produced by the weaker ligand to that produced by the stronger ligand, and the ligand interference coefficient (LIC), which measures the degree to which two ligands positively or negatively synergize. We computed these values for each of the 16 responses of the cell type library for each set of biochemical parameters and determined what response classes each fell into, adapting previously described criteria (Antebi et al., 2017). Ratiometric responses were defined by RLS < 0.2, additive responses by RLS > 0.8 and |LIC| < 0.05, imbalance responses by RLS > 0.8 and LIC < −0.1, and balance responses by RLS > 0.8 and LIC > 0.1. Responses outside these ranges were considered to be intermediate variants and not classified as a specific archetype.

##### Relationship with Mutual Information

Having identified the response classes represented for each set of biochemical parameters, we plotted the distribution of mutual information values associated with a given number of response classes (Figure 6F).

#### Analysis of Parameter Correlations

Based on observations from parameter sets with high mutual information, we computed two correlation measures for the biochemical parameters. Since each parameter could only take on two values, we transformed them to −1 and 1. In particular, we defined 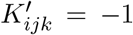 if *K_ijk_* = 0.1 (low) and 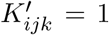 if *K_ijk_* = 1 (high), with 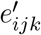 defined analogously. (Equivalently, we defined 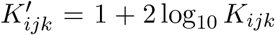) We computed 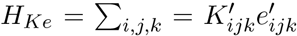 to measure correlation between binding and signaling efficiency for each signaling complex. We also computed 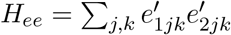, measuring the correlation between activity of the two signaling complexes with the same receptor dimer but different ligands.

We calculated these correlation metrics for each of the 65,536 parameter sets from systematic lattice sampling. To investigate the relationship with mutual information, we sorted parameter sets by mutual information, binned them (with a bin size of 800), and computed the mean correlation and mutual information in each bin (Figures 6G-H).

#### Evolutionary Algorithm as a Generative Model

While the observed anticorrelations of *K* and *e* appear to be predictive of mutual information, it is not clear if these relationships are sufficient to fully describe the criteria for high addressing power and can thus serve as a design principle. Therefore, we developed a generative algorithm that systematically evolves a given set of parameters *K, e* according to these principles and asked whether favoring anticorrelations yields higher addressing power.

We first formed a fitness function *F* (*K, e*) = *−* (*H_Ke_* + *H_ee_*), where *H_Ke_* and *H_ee_* are as above. Intuitively, a choice of *K, e* that has strong affinity-activity or activity-activity anticorrelations would have high fitness. Our algorithm is then a simple “evolutionary” algorithm that performs noisy gradient ascent in this fitness landscape. Starting with a given *K, e*, each iteration involves choosing a random element of *K, e* and proposing a flip (changing it to high if currently low or vice versa). We then compute the resulting change in fitness Δ*F*. If such a flip increases fitness (i.e., Δ*F >* 0), we immediately implement it. If the proposed flip decreases fitness (i.e., Δ*F* < 0), we accept it with a probability *e^sΔF^*, where *s* represents the selection pressure. Such moves towards lower fitness allow dynamics to escape local fitness maxima; the frequency of such moves towards lower fitness is controlled by the selection pressure *s* (or, equivalently, temperature in Monte Carlo algorithms). We repeat this process over many iterations and track the addressing power of the *K, e* configuration at each step.

We first ran our algorithm on 2000 randomly initialized choices of *K, e*. We stop the algorithm after 200 iterations and quantify the addressability of the resulting *K, e* using mutual information. We then visualized the resulting distribution of mutual information values (Figure 6K). With *s* = 0 (i.e., no selection for correlations) produces a wide histogram equivalent to random sampling of parameter space; indeed, only a few parameter sets show significant addressing power. However, with *s* = 1 and therefore selection for parameter anticorrelations, the resulting histogram is significantly shifted towards higher addressing power, despite starting from similar randomly chosen initial conditions. (Longer runs of the evolutionary algorithm did not change the resulting histograms, indicating equilibration within 200 iterations.)

## Notes

http://dx.doi.org/10.22002/D1.1692

### Competing Interest Statement

The authors have declared no competing interest.

